# Prostaglandin EP3 receptor-expressing preoptic neurons bidirectionally control body temperature via tonic GABAergic signaling

**DOI:** 10.1101/2022.04.15.488488

**Authors:** Yoshiko Nakamura, Takaki Yahiro, Naoya Kataoka, Hiroyuki Hioki, Kazuhiro Nakamura

## Abstract

The circuit mechanism of the thermoregulatory center in the preoptic area (POA) is unknown. Using rats, here we show prostaglandin EP3 receptor-expressing POA neurons (POA^EP3R^ neurons) as a pivotal bidirectional controller in the central thermoregulatory mechanism. POA^EP3R^ neurons are activated in response to elevated ambient temperature, but inhibited by prostaglandin E_2_, a pyrogenic mediator. Chemogenetic stimulation of POA^EP3R^ neurons at room temperature reduces body temperature by enhancing heat dissipation, whereas inhibition of them elicits hyperthermia involving brown fat thermogenesis, mimicking fever. POA^EP3R^ neurons innervate sympathoexcitatory neurons in the dorsomedial hypothalamus (DMH) via tonic inhibitory signaling. Although many POA^EP3R^ neuronal cell bodies express a glutamatergic mRNA marker, paradoxically, their axons in the DMH predominantly contain terminals with GABAergic presynaptic proteins, which are increased by chronic heat exposure. These findings indicate that tonic GABAergic inhibitory signaling from POA^EP3R^ neurons is a fundamental determinant of body temperature for thermal homeostasis and fever.

## INTRODUCTION

Thermoregulation is a physiological function fundamental to homeostasis in mammals. Body core temperature is maintained within a control range by autonomous regulation of the balance between heat production within the body and heat loss to the environment. Autonomic thermoregulatory responses, such as sympathetic thermogenesis in brown adipose tissue (BAT) and heat loss control through skin vasomotion, are governed by the CNS. The central regulation of the autonomic thermoregulatory responses is mediated by the thermoregulatory center in the preoptic area (POA) of the hypothalamus and the sympathoexcitatory efferent pathways involving the dorsomedial hypothalamus (DMH) and rostral medullary raphe region (rMR) (Nakamura, 2011; Tan and Knight, 2018; Morrison and Nakamura, 2019; Nakamura et al., 2022). However, the principle of the mechanism by which the POA thermoregulatory center controls the sympathoexcitatory pathways remains to be elucidated.

The POA receives and integrates thermosensory (cool- and warm-sensory) neural signals from skin thermoreceptors and a pyrogenic humoral signal mediated by prostaglandin E_2_ (PGE_2_), which is produced in response to infections (Yamagata et al., 2001; Nakamura and Morrison, 2008, 2010). The POA has been postulated to provide strong tonic inhibitory influences to sympathoexcitatory neurons in the DMH and rMR to regulate the tones of sympathetic outflows to thermoregulatory effectors, because a knife cut to disrupt descending fibers from the POA causes dysregulated increases in BAT thermogenesis and cutaneous vasoconstrictor activity leading to lethal hyperthermia (Chen et al., 1998; Rathner et al., 2008). These findings suggest that the descending tonic inhibition from the POA, whose intensity is altered by thermosensory and pyrogenic afferent signals, determines the basal tone of the sympathoexcitatory efferent signaling to thermoregulatory effector organs. However, the POA neurons that provide such descending tonic inhibitory signaling to bidirectionally control body temperature have yet to be identified.

A group of POA neurons that express the EP3 subtype of prostaglandin E receptors (EP3R) (POA^EP3R^ neurons) has been hypothesized to provide the tonic inhibition (Nakamura et al., 2002; Nakamura, 2011; Morrison and Nakamura, 2019), as supported by the anatomical observations that the POA^EP3R^ neuron group includes GABAergic inhibitory neurons and projects to the DMH and rMR (Nakamura et al., 2002, 2005, 2009). EP3R proteins are localized in neuronal cell bodies and dendritic fibers in the median preoptic nucleus (MnPO) and medial preoptic area (MPA) (Nakamura et al., 1999, 2000) (Figure S1A) and mediate the febrile action of PGE_2_ (Lazarus et al., 2007). However, there is no evidence that POA^EP3R^ neurons are involved in basal thermoregulation. Moreover, the circuit mechanism by which the PGE_2_–EP3R signaling in the POA alters the sympathetic outflows through the DMH and rMR to develop fever remains to be determined. It is even controversial whether the POA^EP3R^ neuron group responsible for fever development is GABAergic or glutamatergic (Machado et al., 2020; Nakamura et al., 2022).

To address these issues, in this study, we investigated the physiological role of POA^EP3R^ neurons in the central circuit mechanisms for thermoregulation and fever, by combining *in vivo* physiological, chemogenetic and neuroanatomical approaches with an antibody to EP3R proteins and a newly developed transgenic rat line that expresses the tetracycline-controlled transactivator (tTA) in EP3R-expressing cells (*Ptger3*-tTA rats). We first examined activation of POA^EP3R^ neurons in response to ambient thermal challenges and then, neurotransmitter markers contained by their cell bodies and axon terminals. The impact of chronic heat exposure of animals on the transmitter phenotypes of POA^EP3R^ neurons was also investigated in relevance to the mechanism of heat tolerance. Finally, we examined the effects of chemogenetic manipulations of POA^EP3R^ neuronal activities on sympathetic outflows and thermoregulatory functions.

## RESULTS

### POA^EP3R^ neurons are activated by ambient heat exposure and project to DMH

To investigate whether POA^EP3R^ neurons are activated in response to ambient thermal challenges, we exposed rats to an environment of 4°C (cold exposure), 24°C (control exposure) or 36°C (heat exposure) for 2 hrs and subsequently examined expression of Fos, a marker for neuronal activation, in POA^EP3R^ neurons by double immunofluorescence histochemistry. Following control exposure, only a small number of POA^EP3R^ neurons exhibited Fos expression (35 ± 15 in 1280 ± 492 POA^EP3R^ neurons, *n* = 4 rats; Figures 1A and 1B). Heat exposure remarkably increased Fos-expressing POA^EP3R^ neurons (246 ± 36 in 1525 ± 353 POA^EP3R^ neurons, *n* = 4 rats), but cold exposure did not (36 ± 10 in 1612 ± 299 POA^EP3R^ neurons, *n* = 4 rats; Figures 1A and 1B). Such warming-activated POA^EP3R^ neurons were distributed both in the MnPO and MPA (Figures 1C and S1B). These observations indicate that the POA^EP3R^ neuronal group includes a substantial subpopulation of warming-activated neurons, but not cooling-activated neurons.

**Figure 1.**
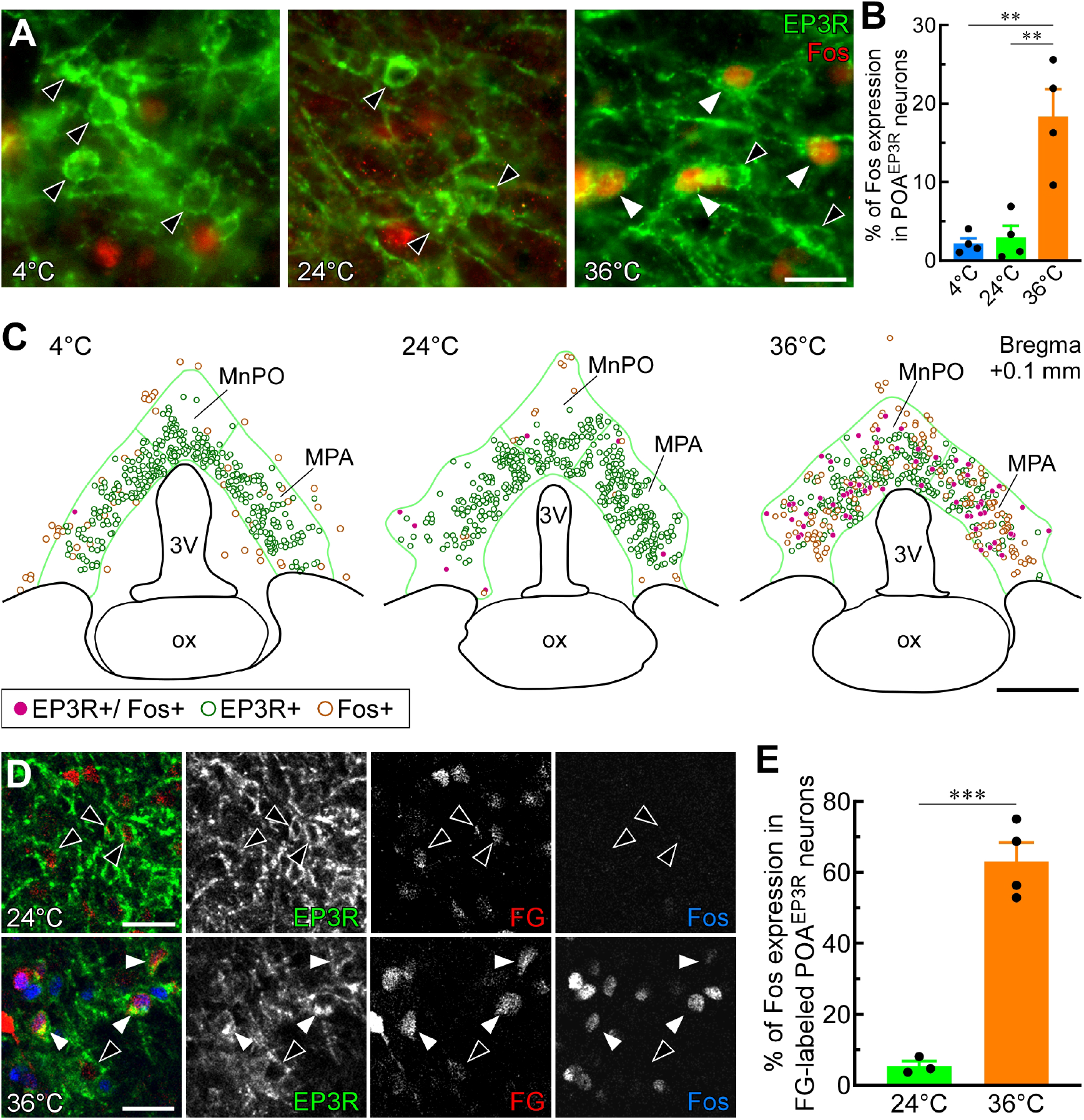
POA^EP3R^ neurons are activated by ambient heat exposure and project to DMH. (A) Double immunofluorescence staining for EP3R and Fos in the POA following 2 hr exposure of rats to ambient temperature of 4°C (cold), 24°C (control) or 36°C (heat). Solid and hollow arrowheads indicate POA^EP3R^ neuronal cell bodies with and without Fos immunoreactivity, respectively. Scale bar, 20 µm. (B) Percentages of Fos expression in POA^EP3R^ neuronal cell bodies (*n* = 4 per group). \*\**P* < 0.01 (Bonferroni’s *post hoc* test following ordinary one-way ANOVA: *F*_2,9_ = 17.05, *P* < 0.001). All values are means ± SEM. (C) Representative distributions of POA^EP3R^ neurons with and without Fos immunoreactivity following exposure to respective temperature. Scale bar, 0.5 mm. 3V, third ventricle; MnPO, median preoptic nucleus; MPA, medial preoptic area, ox, optic chiasm. Green lines delineate EP3R-immunoreactive areas in the MnPO and MPA, which are anatomically defined in Figure S1A. Distributions of cells with EP3R immunoreactivity and/or Fos immunoreactivity in other POA sections are shown in Figure S1B. (D) Pseudocolored confocal images of Fos immunoreactivity in FG-labeled POA^EP3R^ neuronal cell bodies following 2 hr exposure of rats to 24°C or 36°C ambient temperature. Solid and hollow arrowheads indicate FG-labeled POA^EP3R^ neuronal cell bodies with and without Fos immunoreactivity, respectively. Scale bars, 30 µm. Bilateral FG injections in the DMH for all rats were mapped in Figure S1C. Representative distributions of POA cells labeled with FG, EP3R immunoreactivity and/or Fos immunoreactivity are shown in Figure S1D. (E) Percentages of Fos expression in FG-labeled POA^EP3R^ neuronal cell bodies (*n* = 3 for 24°C, *n* = 4 for 36°C). \*\*\**P* < 0.001 (unpaired *t*-test; *t*_5_ = 9.26). All values are means ± SEM.

POA^EP3R^ neurons innervate the DMH, which consists of the dorsomedial hypothalamic nucleus and dorsal hypothalamic area (Nakamura et al., 2005, 2009). To examine whether the warm-responsive POA^EP3R^ neurons project to the DMH, we performed retrograde neural tracing from the DMH. Fluoro-Gold (FG), a retrograde tracer, was bilaterally injected into the hypothalamus to fill the most part of the DMH (Figure S1C) and then, the rats were exposed to 24°C or 36°C for 2 hrs. Heat exposure induced Fos expression in more than 60% of FG-labeled POA^EP3R^ neurons (152 ± 17 in 249 ± 42 FG-labeled POA^EP3R^ neurons, *n* = 4 rats), in contrast to few Fos-positive cells following control exposure (12 ± 2 in 236 ± 35 FG-labeled POA^EP3R^ neurons, *n* = 3 rats) (Figures 1D and 1E). Such triple-labeled neurons were distributed in both MnPO and MPA (Figure S1D). These results indicate that POA^EP3R^→DMH transmission is enhanced under conditions in which heat defense is required for thermal homeostasis.

### PGE_2_ inhibits heat exposure-induced activation of POA^EP3R^ neurons

The EP3R has been shown to be coupled mostly with the inhibitory G protein (Gi) to reduce the intracellular cAMP level in culture cells (Narumiya et al., 1999). Therefore, we questioned whether a PGE_2_ action on POA^EP3R^ neurons inhibits their elevated activity during heat exposure to trigger fever. PGE_2_ or saline was injected into the lateral ventricle 10 min before the rats underwent heat exposure (36°C). PGE_2_ injection elicited an intense fever, which was characterized by a rapid elevation of body core temperature (*T*_core_) by 3.1 ± 0.3°C (*n* = 3) and maintained throughout heat exposure (Figure 2A). Rats exposed to heat following saline injection exhibited Fos expression in approximately 30% of POA^EP3R^ neurons (Figures 2B and 2C). This heat exposure-induced Fos expression in POA^EP3R^ neurons was significantly reduced by PGE_2_ injection (Figures 2B and 2C). A linear regression analysis revealed a significant negative correlation between Fos expression in POA^EP3R^ neurons and *T*_core_ at 60 min of heat exposure (Figure 2D). This analysis predicted that *T*_core_ would be elevated up to 42.1°C if all POA^EP3R^ neurons were suppressed during heat exposure and that the activation level at which approximately 40% of POA^EP3R^ neurons express Fos would be sufficient for the maintenance of *T*_core_ around thermoneutral 37°C during heat exposure (Figure 2D). Although PGE_2_ injection significantly reduced Fos expression in EP3R-expressing neurons in both MnPO and MPA (Figures 2E and S2), there was a tendency toward stronger inhibition in the MnPO (Figure 2E). These results demonstrate that PGE_2_ inhibits heat exposure-induced activation of POA^EP3R^ neurons, consistent with the view that the activity of POA^EP3R^ neurons is dynamically changed by warm-sensory inputs to defend *T*_core_ and by an action of PGE_2_ to trigger fever.

**Figure 2.**
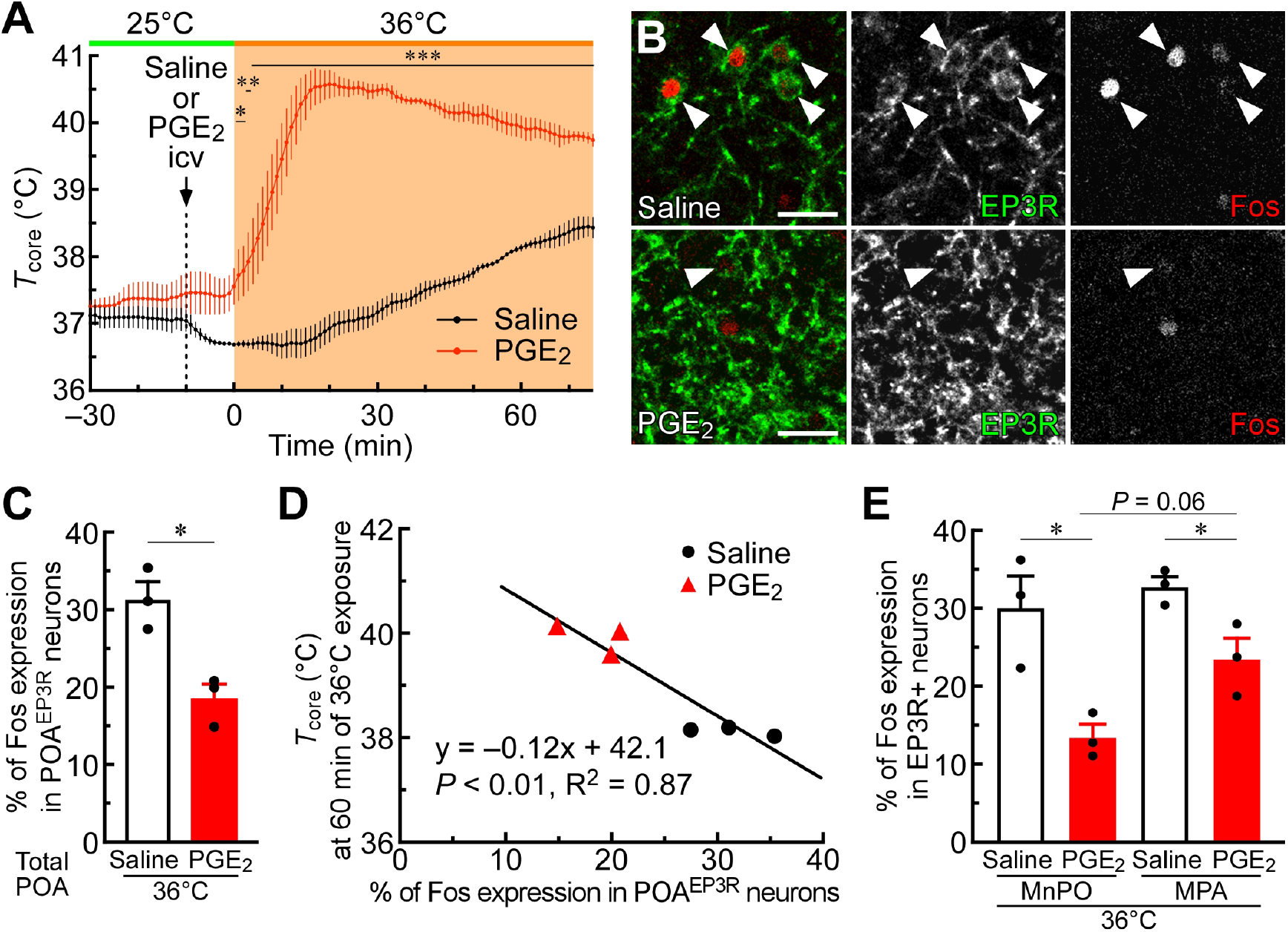
PGE_2_ inhibits heat exposure-induced activation of POA^EP3R^ neurons. (A) Changes in *T*_core_ of rats during exposure to 36°C ambient temperature following intracerebroventricular (icv) injection of saline or PGE_2_ (*n* = 3 per group) were analyzed by repeated measures two-way ANOVA (injectant: *F*_1,4_ = 207.1, *P* < 0.001; time: *F*_105,420_ = 42.74, *P* < 0.001; interaction: *F*_105,420_ = 22.83, *P* < 0.001) followed by Bonferroni’s *post hoc* test (\**P* < 0.05; \*\**P* < 0.01; \*\*\**P* < 0.001, horizontal bars with asterisks indicate time points with significant difference). All values are means ± SEM. (B) Confocal images of Fos immunoreactivity in POA^EP3R^ neuronal cell bodies following icv injection of saline or PGE_2_ and exposure to 36°C ambient temperature. Heat exposure increased Fos-expressing POA^EP3R^ neurons (arrowheads), which were reduced by PGE_2_ injection. Scale bars, 30 µm. Representative distributions of POA cells with EP3R immunoreactivity and/or Fos immunoreactivity are shown in Figure S2. (C) Percentages of Fos expression in POA^EP3R^ neurons following heat exposure after saline or PGE_2_ injection (*n* = 3 per group). \**P* < 0.05 (unpaired *t*-test; *t*_4_ = 4.34). All values are means ± SEM. (D) Relationship between the percentage of Fos expression in POA^EP3R^ neurons and *T*_core_ of the rats at 60 min of heat exposure. Data from the saline-injected and PGE_2_- injected groups were subjected to linear regression analysis (Pearson’s correlation test). (E) Percentages of Fos expression in EP3R-immunoreactive cell bodies in the MnPO and MPA following heat exposure after saline or PGE_2_ injection (*n* = 3 per group). \**P* < 0.05 (unpaired *t*-tests; Saline vs PGE_2_ in MnPO: *t*_4_ = 3.78; Saline vs PGE_2_ in MPA: *t*_4_ = 3.12; paired *t*-tests; MnPO-Saline vs MPA-Saline: *t*_2_ = 0.67; MnPO-PGE_2_ vs MPA-PGE_2_: *t*_2_ = 3.83). All values are means ± SEM.

### POA^EP3R^ neuronal cell bodies predominantly express glutamatergic marker rather than GABAergic marker

We next investigated the transmitter phenotype of POA^EP3R^ neurons. *In situ* hybridization for mRNA of glutamic acid decarboxylase 1 (GAD67, also known as GAD1), a marker for GABAergic neurons, or vesicular glutamate transporter 2 (VGLUT2), a marker for glutamatergic neurons, was performed in combination with EP3R immunohistochemistry in the POA. GAD67 and VGLUT2 mRNAs were detected in 22% and 44% of POA^EP3R^ neuronal cell bodies, respectively (GAD67: 352 ± 12 in 1,676 ± 198 POA^EP3R^ cell bodies; VGLUT2: 559 ± 55 in 1,299 ± 168 POA^EP3R^ cell bodies, *n* = 4 rats; Figures 3A and 3B). The percentage of POA^EP3R^ cell bodies expressing GAD67 mRNA was lower than that obtained from our previous double-labeling analysis with bright-field microscopy (86%; Nakamura et al., 2002), which perhaps made it easier to pick up false-positive cells. Whereas GAD67 and VGLUT2 mRNA expressions were detected in similar percentages of POA^EP3R^ neuronal cell bodies in the rostral half of the POA, the VGLUT2-expressing population was increased in the caudal half of the POA (Figures S3A and S3B).

**Figure 3.**
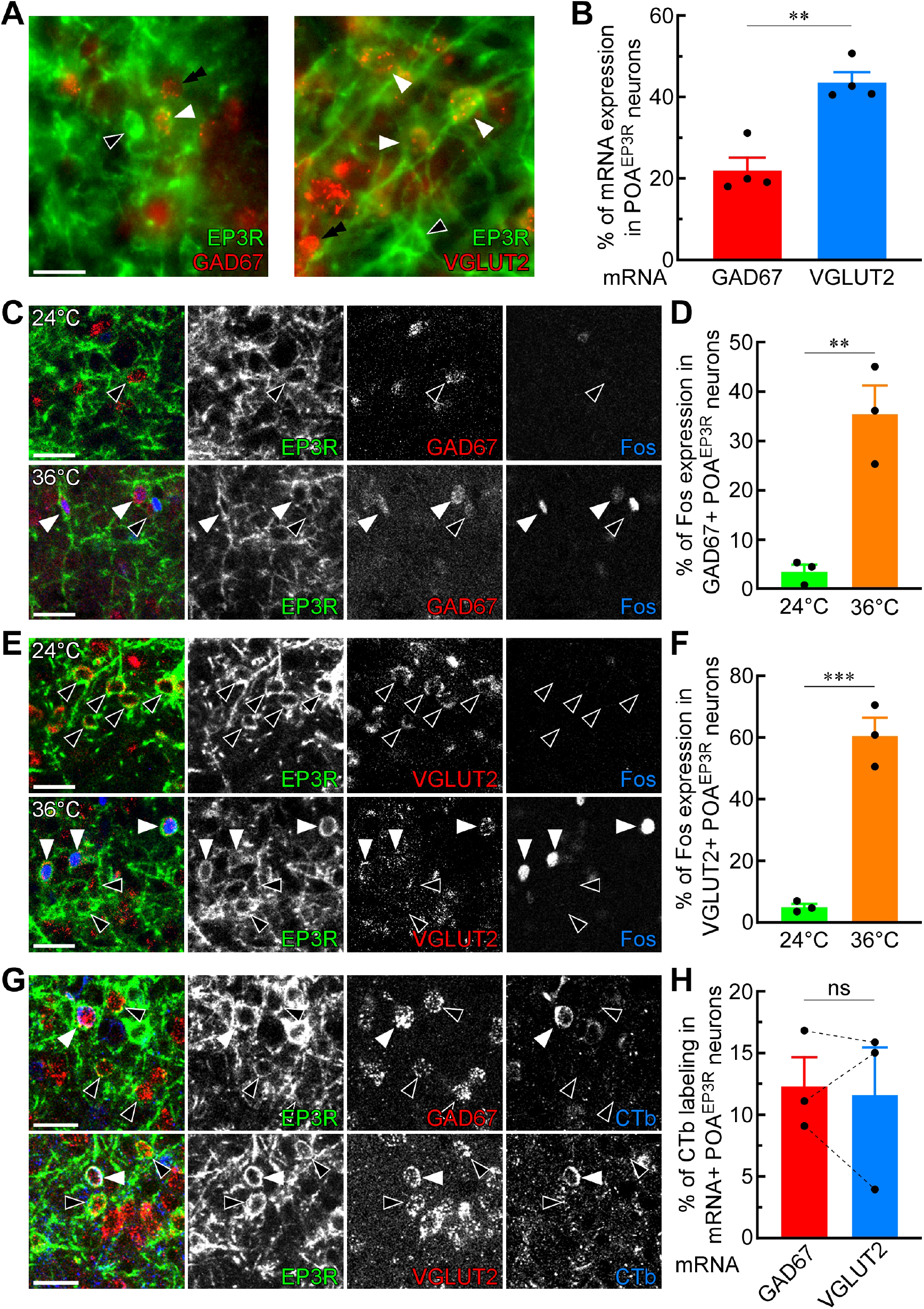
POA^EP3R^ neuronal cell bodies predominantly express glutamatergic marker rather than GABAergic marker. (A) Double fluorescence labeling with EP3R immunoreactivity and GAD67 or VGLUT2 mRNA hybridization signals in POA neurons. Solid and hollow arrowheads indicate POA^EP3R^ neuronal cell bodies with and without mRNA signals, respectively. Double arrowheads indicate mRNA signals in EP3R-immunonegative cells. Scale bar, 20 µm. Representative distributions of POA^EP3R^ neurons with and without GAD67 or VGLUT2 mRNA signals are shown in Figures S3A and S3B. (B) Percentages of GAD67 and VGLUT2 mRNA expression in POA^EP3R^ neuronal cell bodies (*n* = 4 per group). \*\**P* < 0.01 (unpaired *t*-test; *t*_6_ = 5.56). All values are means ± SEM. (C–F) Confocal images of Fos immunoreactivity in GAD67 mRNA-expressing (C) or VGLUT2 mRNA-expressing (E) POA^EP3R^ neuronal cell bodies following 2 hr exposure of rats to 24°C or 36°C ambient temperature. Solid and hollow arrowheads indicate mRNA-expressing POA^EP3R^ neuronal cell bodies with and without Fos immunoreactivity, respectively. Scale bars, 30 µm. Exposure to 36°C, compared to 24°C, increased Fos expression in POA^EP3R^ neurons expressing GAD67 (D) and VGLUT2 (F) mRNA (*n* = 3 per group). \*\**P* < 0.01; \*\*\**P* < 0.001 [unpaired *t*-tests; *t*_4_ = 5.43 (D), *t*_4_ = 9.50 (F)]. All values are means ± SEM. Representative distributions of POA cells with EP3R immunoreactivity, Fos immunoreactivity and/or GAD67 or VGLUT2 mRNA hybridization signals are shown in Figures S3C and S3D. (G and H) Pseudocolored confocal images of GAD67 (top, G) or VGLUT2 (bottom, G) mRNA-expressing POA^EP3R^ neuronal cell bodies that exhibited immunoreactivity for CTb derived from the DMH. Solid and hollow arrowheads indicate mRNA-expressing POA^EP3R^ neuronal cell bodies with and without CTb immunoreactivity, respectively. Scale bars, 30 µm. Percentages of CTb labeling were comparable between GAD67 mRNA-expressing and VGLUT2 mRNA-expressing populations of POA^EP3R^ neurons (H) (*n* = 3). ns, not significant (paired *t*-test; *t*_2_ = 0.27). All values are means ± SEM. Sites of unilateral CTb injections in the DMH for all the rats are shown in Figures S3E and S3F. Percentages of GAD67 and VGLUT2 mRNA expression in CTb-labeled POA^EP3R^ neuronal cell bodies are shown in Figure S3G.

To examine whether POA^EP3R^ neurons expressing GAD67 or VGLUT2 mRNA were activated by heat exposure, we performed triple fluorescence labeling combining EP3R and Fos immunohistochemistry and GAD67 or VGLUT2 *in situ* hybridization in the POA of rats exposed to 24°C or 36°C for 2 hrs. Consistent with the aforementioned results, Fos expression following control exposure was detected in only 3∼5% of POA^EP3R^ cell bodies expressing GAD67 or VGLUT2 mRNA (Figures 3C– 3F). In contrast, heat exposure induced Fos expression in 36% and 61% of POA^EP3R^ cell bodies expressing GAD67 or VGLUT2 mRNA, respectively (72 ± 24 in 193 ± 36 GAD67-expressing POA^EP3R^ cell bodies; 791 ± 63 in 1343 ± 227 VGLUT2-expressing POA^EP3R^ cell bodies, *n* = 3 rats; Figures 3C–3F). Heat exposure-induced Fos expression in POA^EP3R^ cell bodies expressing either mRNA was observed in both MnPO and MPA (Figures S3C and S3D). Therefore, both GAD67 mRNA-expressing and VGLUT2 mRNA-expressing groups of POA^EP3R^ neurons are activated in response to elevation of ambient temperature.

We further performed retrograde tracing to determine whether POA^EP3R^ neurons expressing GAD67 or VGLUT2 mRNA projected to the DMH. Fluorophore-conjugated cholera toxin b-subunit (CTb), a retrograde tracer, was unilaterally injected to cover a part of the DMH (Figures S3E and S3F). With this size of injection, CTb was detected in 12% of either group of POA^EP3R^ cell bodies with GAD67 or VGLUT2 mRNA (16 ± 2 in 138 ± 30 GAD67-expressing POA^EP3R^ cells; 85 ± 38 in 695 ± 162 VGLUT2-expressing POA^EP3R^ cells, *n* = 3 rats; Figures 3G and 3H). CTb-labeled POA^EP3R^ neurons consisted of 20% of GAD67-expressing cells and 81% of VGLUT2-expressing cells (Figure S3G). These results indicate that both GAD67 mRNA-and VGLUT2 mRNA-expressing groups of POA^EP3R^ neurons provide axonal projections to the DMH.

### POA^EP3R^→DMH axon terminals predominantly contain GABAergic marker rather than glutamatergic marker

To more directly determine the neurotransmitters used for POA^EP3R^→DMH transmission, we sought to perform anterograde tract tracing specifically from POA^EP3R^ neurons. For this purpose, we generated the *Ptger3*-tTA transgenic rat line in which tTA was expressed under the promoter of the *Ptger3* (EP3R) gene (Figure S4). Injection into the POA of *Ptger3*-tTA rats with the adeno-associated virus (AAV) vector, AAV-TRE-palGFP, to express palGFP, a membrane-targeted form of GFP (Moriyoshi et al., 1996), under the control of the tetracycline response element (TRE) resulted in highly selective labeling of POA^EP3R^ neurons with palGFP: distribution of palGFP-labeled cells well overlapped with that of POA^EP3R^ neurons (Figures 4A and 4B) and 97% of palGFP-labeled neurons were immunopositive for EP3R.

**Figure 4.**
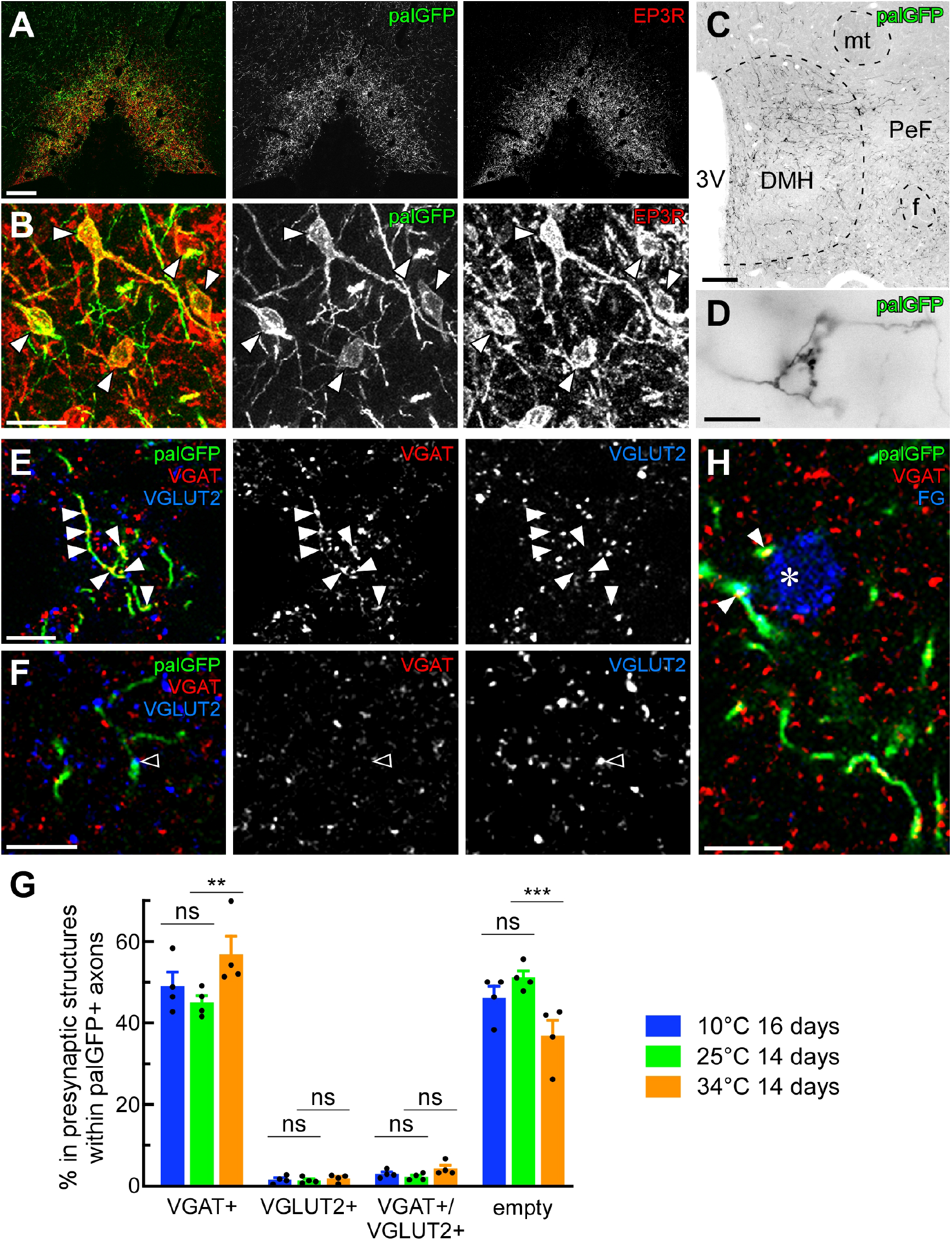
POA^EP3R^→DMH axon terminals predominantly contain GABAergic marker rather than glutamatergic marker. (A and B) Selective transduction of POA^EP3R^ neurons with palGFP by injecting AAV-TRE-palGFP into the POA of *Ptger3*-tTA rats (A). Magnified confocal images (B) show palGFP-expressing POA^EP3R^ neurons (arrows). Scale bars, 0.2 mm (A) and 30 µm (B). See also Figure S4 for the production of *Ptger3*-tTA rats. (C and D) The composite fluorescence image (C) shows the distribution of POA^EP3R^ neuron-derived, palGFP-immunoreactive axons in the DMH and perifornical area (PeF). The magnified image (D) shows palGFP-labeled basket-like structures with axonal boutons in the DMH. Scale bars, 0.2 mm (C) and 10 µm (D). f, fornix; mt, mammillothalamic tract. Distributions of POA^EP3R^ neuron-derived, palGFP-labeled axons in other brain regions are shown in Figure S5. (E and F) Confocal images of palGFP-labeled (POA^EP3R^ neuron-derived) axons with VGAT-immunoreactive puncta (solid arrowheads in E) and VGLUT2-immunoreactive puncta (hollow arrowhead in F) in the DMH. Scale bars, 10 µm. Other confocal images of presynaptic structures in the DMH are presented in Figure S6. (G) Percentages of VGAT-immunoreactive puncta (VGAT+), VGLUT2-immunoreactive puncta (VGLUT2+), double immunoreactive puncta (VGAT+/VGLUT2+) and double immunonegative axon swellings (empty) in total of the axonal presynaptic structures in palGFP-labeled axons in the DMH of *Ptger3*-tTA rats exposed to 10°C, 25°C or 34°C for 14–16 days (*n* = 4 per group). Data were analyzed by repeated measures two-way ANOVA (transmitter marker: *F*_3,27_ = 309.8, *P* < 0.001; temperature: *F*_2,9_ = 0.43, *P* = 0.66; interaction: *F*_6,27_ = 4.52, *P* < 0.01) followed by Bonferroni’s *post hoc* test (ns, not significant; \*\**P* < 0.01; \*\*\**P* < 0.001). All values are means ± SEM. (H) Pseudocolored confocal image of VGAT-immunoreactive, palGFP-labeled (POA^EP3R^ neuron-derived) axon swellings (arrowheads) that are apposed to a DMH neuronal cell body (asterisk) retrogradely labeled with FG from the rMR. Scale bar, 10 µm. A representative injection of FG into the rMR and the resultant retrograde labeling of DMH neurons with FG are shown in Figures S7C and S7D.

The expression of palGFP clearly visualized axonal morphology of POA^EP3R^ neurons to their terminals, and palGFP-labeled axons were widely observed in the hypothalamus, thalamus, amygdala, midbrain, pons and medulla oblongata. Dense distribution of palGFP-labeled axon fibers and terminals was observed in the DMH (Figure 4C). Other brain regions in which palGFP-labeled axon fibers and terminals were distributed include the paraventricular hypothalamic nucleus, hypothalamic arcuate nucleus, perifornical area, medial tuberal nucleus, peduncular part of the lateral hypothalamus, anterior parts of the mediodorsal and paraventricular thalamic nuclei, lateral habenular nucleus, basolateral amygdaloid nucleus, ventral subiculum, lateral and ventrolateral periaqueductal gray, lateral parabrachial nucleus, raphe magnus nucleus, rostral raphe pallidus nucleus, and a transition area between the caudal raphe pallidus and raphe obscurus nuclei (Figures 4C and S5).

The palGFP-labeled axons in the DMH often formed basket-like structures with axonal boutons (Figure 4D), putative synaptic contacts. Therefore, we performed confocal laser-scanning microscopy to detect vesicular transporters for the fast transmitters, GABA and glutamate, in palGFP-labeled DMH-projecting axons of POA^EP3R^ neurons. Unexpectedly, palGFP-labeled axons in the DMH contained abundant puncta immunoreactive for vesicular GABA transporter (VGAT), a marker for GABAergic presynaptic terminals, whereas VGLUT2 immunoreactive puncta, indicating glutamatergic terminals, were only occasionally found in these axons (Figures 4E, 4F and S6A; for quantification data, see below and Figure 4G). Some of the VGLUT2-immunoreactive puncta also exhibited VGAT immunoreactivity (Figure S6B).

Most VGAT-immunoreactive puncta in palGFP-labeled axons in the DMH were immunoreactive for synaptophysin (Figure S6C), confirming that these puncta constitute presynaptic structures. Also, immunoreactivity for GAD67, a GABA-synthesizing enzyme, was colocalized with or in close proximity to many of VGAT-immunoreactive puncta in palGFP-labeled axons in the DMH (Figure S6D), indicating the capability of local GABA production in those VGAT-containing presynaptic structures. Validating the specificity of the anti-VGAT guinea pig antibody used for the histochemical analyses, VGAT-immunoreactive profiles visualized by this antibody in the DMH were co-labeled well with mouse and rabbit antibodies raised against different epitopes on the VGAT protein (Figure S6E). To validate our VGLUT2 immunohistochemistry, we tested whether the anti-VGLUT2 antibody used can visualize axon terminals of thalamocortical neurons, which mostly express VGLUT2 (Kaneko and Fujiyama, 2002). Injection of AAV-TRE-palGFP into the anterior midline thalamus of *Ptger3*-tTA rats transduced many neurons in the anterior part of the paraventricular thalamic nucleus and central medial thalamic nucleus with palGFP (Figure S7A), consistent with intense EP3R expression in these nuclei (Nakamura et al., 2000). Their palGFP-labeled axons were densely distributed in the area 2 of the cingulate cortex and contained many VGLUT2-immunoreactive puncta (Figure S7B), but few VGAT-immunoreactive puncta (83% and 4% of 1,725 immunoreactive puncta counted in palGFP-labeled axons in the area 2 of the cingulate cortex, respectively). Some double immunoreactive puncta (13%) were also observed. These data validate our immunohistochemical analyses of transporter localization in POA^EP3R^→DMH axon terminals.

The DMH contains glutamatergic neurons that transmit sympathoexcitatory signals to sympathetic premotor neurons in the rMR to drive thermogenic responses for cold defense and fever (Zaretskaia et al., 2003; Madden and Morrison, 2004; Nakamura et al., 2004; Nakamura et al., 2005; Nakamura and Morrison, 2007, 2011; Kataoka et al., 2014). Retrograde labeling with FG injected into the rMR visualized a dense cluster of rMR-projecting neurons in the DMH (Figures S7C and S7D; Samuels et al., 2004; Nakamura et al., 2005). These FG-labeled cell bodies were often closely associated with palGFP-labeled axon swellings (168 ± 20 swellings in contact were found, *n* = 4 rats), suggestive of synaptic contacts, and a substantial number (60 ± 6) of these axon swellings contained VGAT (Figure 4H), but few (3 ± 2) were VGLUT2-positive (additionally, 3 ± 2 swellings in contact were immunoreactive for both VGAT and VGLUT2). These observations support the view that POA^EP3R^→DMH projection neurons regulate DMH→rMR sympathoexcitatory neurons via GABAergic synaptic inputs rather than glutamatergic inputs.

### Chronic heat exposure increases GABAergic POA^EP3R^→DMH axon terminals

Because POA^EP3R^ neurons are responsive to ambient thermal challenges (Figure 1), we questioned whether the localizations of the vesicular transporters in POA^EP3R^→DMH axon terminals were altered by chronic exposure to extreme temperatures. *Ptger3*-tTA rats injected with AAV-TRE-palGFP into the POA were exposed to 10°C (cold exposure), 25°C (control exposure) or 34°C (heat exposure) for 14–16 days, and we subsequently quantified VGAT- and VGLUT2-immunopositive puncta in palGFP-labeled axons in the DMH as presynaptic sites that release GABA and glutamate, respectively. Because there were also a substantial number of palGFP-labeled axon swellings that contained neither VGAT-nor VGLUT2-immunoreactive profile, we also quantified such ‘empty’ swellings that were more than 1.5 times thicker than the adjacent axon shaft; most of swellings that exceeded this threshold thickness were found to contain synaptophysin in our preliminary analyses.

Among 6,123 ± 1,152 of palGFP-labeled axonal presynaptic structures (i.e., total of VGAT- and/or VGLUT2-immunoreactive puncta and empty swellings) counted bilaterally in the DMH of rats (*n* = 4) exposed to 25°C, 45% were VGAT-immunoreactive puncta and 51% were empty swellings (Figure 4G). In contrast, only 1% were VGLUT2-immunoreactive puncta and 2% were double immunoreactive puncta (Figure 4G). Following chronic heat exposure, the population of VGAT-immunoreactive puncta was significantly increased to 57% and empty swellings were reduced to 37%, whereas the VGLUT2-immunoreactive populations were not affected (8,065 ± 2,432 palGFP-labeled axonal presynaptic structures counted in the DMH; *n* = 4 rats; Figure 4G). Chronic cold exposure did not alter any of the populations (6,462 ± 2,284 palGFP-labeled axonal presynaptic structures counted in the DMH; *n* = 4 rats), compared to control exposure (Figure 4G). These data suggest a mechanism of heat adaptation, in which POA^EP3R^ neurons increase GABAergic synapses onto sympathoexcitatory DMH neurons under chronic heat exposure, so that they can more efficiently inhibit thermogenic and other sympathetic outflows to tolerate heat.

### Chemogenetic manipulations of POA^EP3R^ neurons bidirectionally alter body temperature by eliciting effector responses

We examined the effects of selective *in vivo* inhibition and stimulation of POA^EP3R^ neurons on thermoregulation by employing chemogenetic techniques with hM4Di and hM3Dq, Gi- and Gq-coupled designer receptors exclusively activated by designer drug (DREADDs) (Roth, 2016), respectively (Figure 5). To inhibit POA^EP3R^ neurons, we used hM4Di^nrxn^, an axon-targeted variant of hM4Di conjugated with an intracellular domain of neurexin 1α, whose activation can suppress synaptic release probability (Stachniak et al., 2014). POA^EP3R^ neurons were selectively transduced with hM4Di^nrxn^ by injecting AAV-TRE-mCherry-T2A-hM4Di^nrxn^ into the POA of *Ptger3*-tTA rats (Figures 5A and 5B). In these rats, an injection into the lateral ventricle with DREADD Agonist 21 (C21), a selective actuator for the DREADDs (Thompson et al., 2018), increased *T*_core_ and BAT temperature (*T*_BAT_) by 2.5 ± 0.2°C and 2.6 ± 0.2°C (*n* = 4), respectively, whereas saline injection had no effect on the temperatures (Figure 5D). In the initial phase of the temperature rise, the elevation of *T*_BAT_ led that of *T*_core_ (Figure 5G), suggesting that BAT was a source of heat for the development of the hyperthermia. On the other hand, activity levels of the rats, which could also contribute to heat production, were not obviously affected by C21 (Figure 5D). The hyperthermic state was maintained for approximately 8 hrs and returned to baseline by 9 hrs after C21 injection (Figure 5D).

**Figure 5.**
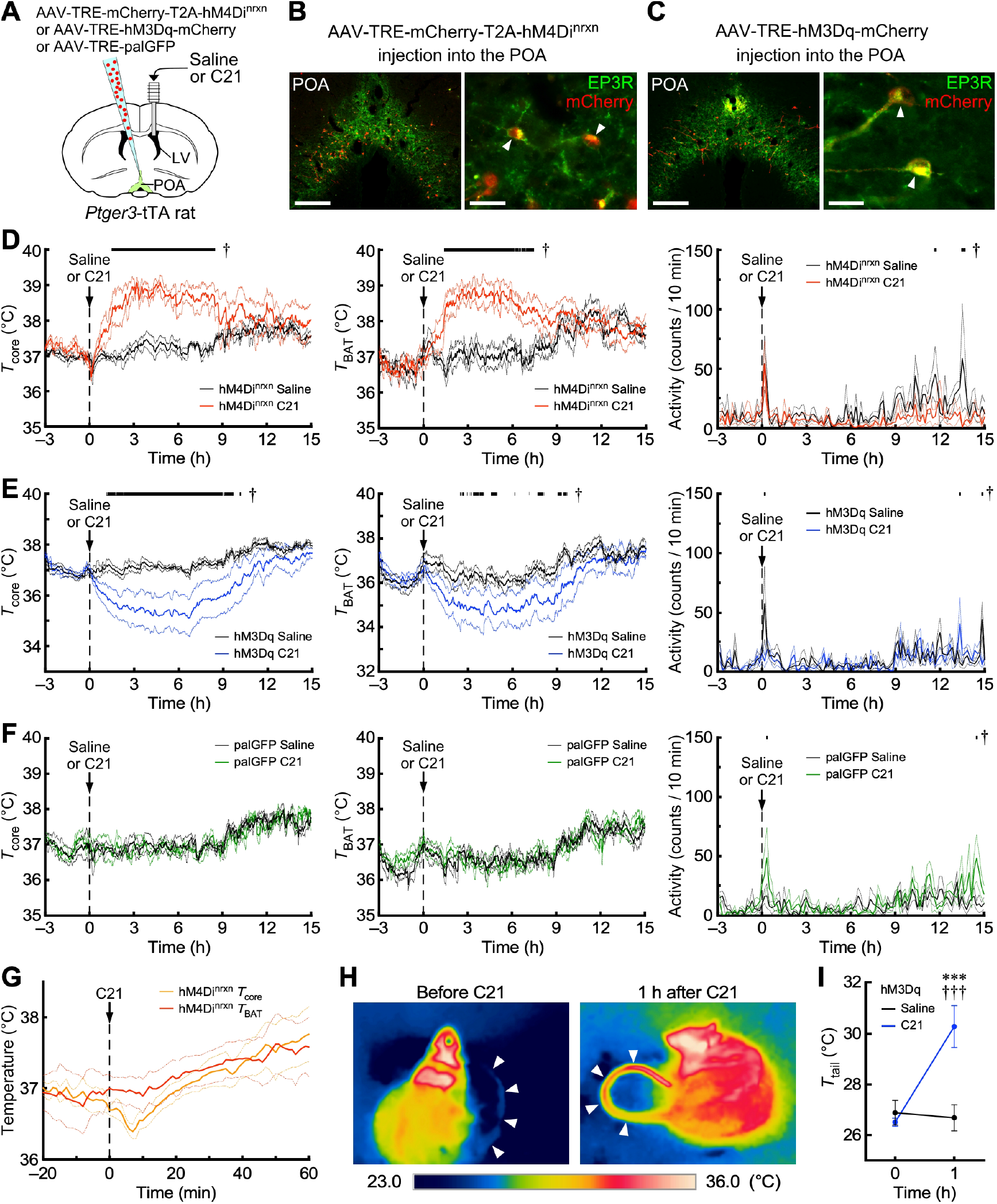
Chemogenetic manipulations of POA^EP3R^ neurons bidirectionally alter body temperature by eliciting effector responses. (A) The injection scheme for *in vivo* chemogenetic experiments. LV, lateral ventricle. (B and C) Representative examples of selective transduction of POA^EP3R^ neurons with mCherry-T2A-hM4Di^nrxn^ (B) and hM3Dq-mCherry (C) in *Ptger3*-tTA rats. Arrowheads indicate POA^EP3R^ neuronal cell bodies labeled with mCherry. Scale bars, 0.3 mm (left panels) and 30 µm (right panels). (D–G) Time-course changes in *T*_core_, *T*_BAT_ and activity following intracerebroventricular injection of saline or C21 in *Ptger3*-tTA rats with POA^EP3R^ neurons transduced with mCherry-T2A-hM4Di^nrxn^ (*n* = 4 per group) (D), hM3Dq-mCherry (*n* = 5 per group) (E) or palGFP (*n* = 4 per group) (F). The injection was made at 10 am and the dark period started at 9 hrs after the injection. Data were analyzed by repeated measures two-way ANOVA [(D) *T*_core_, injectant: *F*_1,3_ = 26.55, *P* = 0.014; time: *F*_1080,3240_ = 3.74, *P* < 0.001; interaction: *F*_1080,3240_ = 4.52, *P* < 0.001; *T*_BAT_, injectant: *F*_1,3_ = 10.65, *P* = 0.047; time: *F*_1080,3240_ = 6.49, *P* < 0.001; interaction: *F*_1080,3240_ = 6.62, *P* < 0.001; activity, injectant: *F*_1,3_ = 13.70, *P* = 0.034; time: *F*_107,321_ = 2.10, *P* < 0.001; interaction: *F*_107,321_ = 1.19, *P* = 0.124; (E) *T*_core_, injectant: *F*_1,4_ = 4.75, *P* = 0.095; time: *F*_1080,4320_ = 10.76, *P* < 0.001; interaction: *F*_1080,4320_ = 4.82, *P* < 0.001; *T*_BAT_, injectant: *F*_1,4_ = 2.67, *P* = 0.178; time: *F*_1080,4320_ = 10.40, *P* < 0.001; interaction: *F*_1080,4320_ = 3.14, *P* < 0.001; activity, injectant: *F*_1,4_ = 5.88, *P* = 0.072; time: *F*_107,428_ = 2.68, *P* < 0.001; interaction: *F*_107,428_ = 1.51, *P* = 0.002; (F) *T*_core_, injectant: *F*_1,3_ = 0.03, *P* = 0.876; time: *F*_1080,3240_ = 8.90, *P* < 0.001; interaction: *F*_1080,3240_ = 0.95, *P* = 0.869; *T*_BAT_, injectant: *F*_1,3_ = 0.39, *P* = 0.577; time: *F*_1080,3240_ = 8.45, *P* < 0.001; interaction: *F*_1080,3240_ = 1.08, *P* = 0.052; activity, injectant: *F*_1,3_ = 2.22, *P* = 0.233; time: *F*_107,321_ = 2.34, *P* < 0.001; interaction: *F*_107,321_ = 1.46, *P* = 0.006] followed by Bonferroni’s *post hoc* test (horizontal bars with † indicate time points with difference at a statistically significant level of *P* < 0.05, *P* < 0.01 or *P* < 0.001). The graph in (G) compares the changes in *T*_core_ and *T*_BAT_ for 60 min after C21 injection in *Ptger3*-tTA rats with POA^EP3R^ neurons transduced with mCherry-T2A-hM4Di^nrxn^ (*n* = 4 per group). All values are means ± SEM. (H and I) Thermographic measurements of the tail skin (arrowheads) in *Ptger3*-tTA rats with POA^EP3R^ neurons transduced with hM3Dq-mCherry (*n* = 4) before and 1 hr after C21 injection. The difference in tail skin temperature (*T*_tail_) at each time point was analyzed by repeated measures two-way ANOVA (injectant: *F*_1,3_ = 17.17, *P* = 0.026; time: *F*_1,3_ = 5.69, *P* = 0.097; interaction: *F*_1,3_ = 405.6, *P* < 0.001) followed by Bonferroni’s *post hoc* test. \*\*\**P* < 0.001 (*vs* saline); ^†††^*P* < 0.001 (*vs* time 0). All values are means ± SEM.

For stimulation of POA^EP3R^ neurons, AAV-TRE-hM3Dq-mCherry was injected into the POA of *Ptger3*-tTA rats to selectively express hM3Dq in POA^EP3R^ neurons (Figures 5A and 5C). In contrast to the hyperthemia induced by inhibition of POA^EP3R^ neurons, injection of C21 into the lateral ventricle of *Ptger3*-tTA rats with POA^EP3R^ neurons expressing hM3Dq induced hypothermia: *T*_core_ and *T*_BAT_ were reduced by 1.9 ± 0.9°C and 2.2 ± 0.9°C (*n* = 5), respectively (Figure 5E). Thermography during the drop in *T*_core_ revealed marked increases in tail skin temperature as well as whole-body surface temperature (Figures 5H and 5I), indicating the induction of an active heat-loss response via increased skin blood flow. The hypothermic *T*_core_ (below 36.0°C) was maintained until 7–9 hrs after C21 injection (Figure 5E). Saline injection induced no obvious change in *T*_core_, *T*_BAT_ or tail skin temperature (Figures 5E and 5I). C21 did not affect activity levels of the rats (Figure 5E). In control *Ptger3*-tTA rats, which expressed palGFP in POA^EP3R^ neurons, C21 had no effect on *T*_core_, *T*_BAT_ or activity (Figure 5F). These *in vivo* physiological data demonstrate that changes in the activity of POA^EP3R^ neurons alter *T*_core_ bidirectionally via altering the regulations of BAT thermogenesis and cutaneous vasoconstrictors.

### POA^EP3R^→DMH transmission tonically inhibits sympathetic outflow to BAT

The present anatomical and physiological results suggest that POA^EP3R^→DMH projection neurons provide tonic GABAergic inhibitory signals to regulate the DMH→rMR sympathoexcitatory pathway, and a reduction of the GABAergic signal tones results in disinhibition of the sympathoexcitatory pathway to drive BAT thermogenesis and hyperthermia. Consistent with this possibility, local inhibition of POA^EP3R^→DMH axon terminals by bilateral nanoinjections of C21 into the DMH of *Ptger3*-tTA rats that expressed hM4Di^nrxn^ in POA^EP3R^ neurons elicited robust activation of BAT thermogenesis and tachycardia in an anesthetized preparation (Figures 6A and 6B). Such obvious sympathetic responses to C21 injections were observed in 2 out of 6 rats we tested. This inconstant observation is likely because the size of the infected population in POA^EP3R^ neurons was variable among animals. In the rats that did not exhibit the C21-induced responses, non-infected POA^EP3R^ neuronal fibers might be dominant and active enough to suppress the sympathoexcitatory DMH neurons under anesthetic conditions even when C21 inhibited hM4Di^nrxn^-containing POA^EP3R^ neuronal axon terminals.

**Figure 6.**
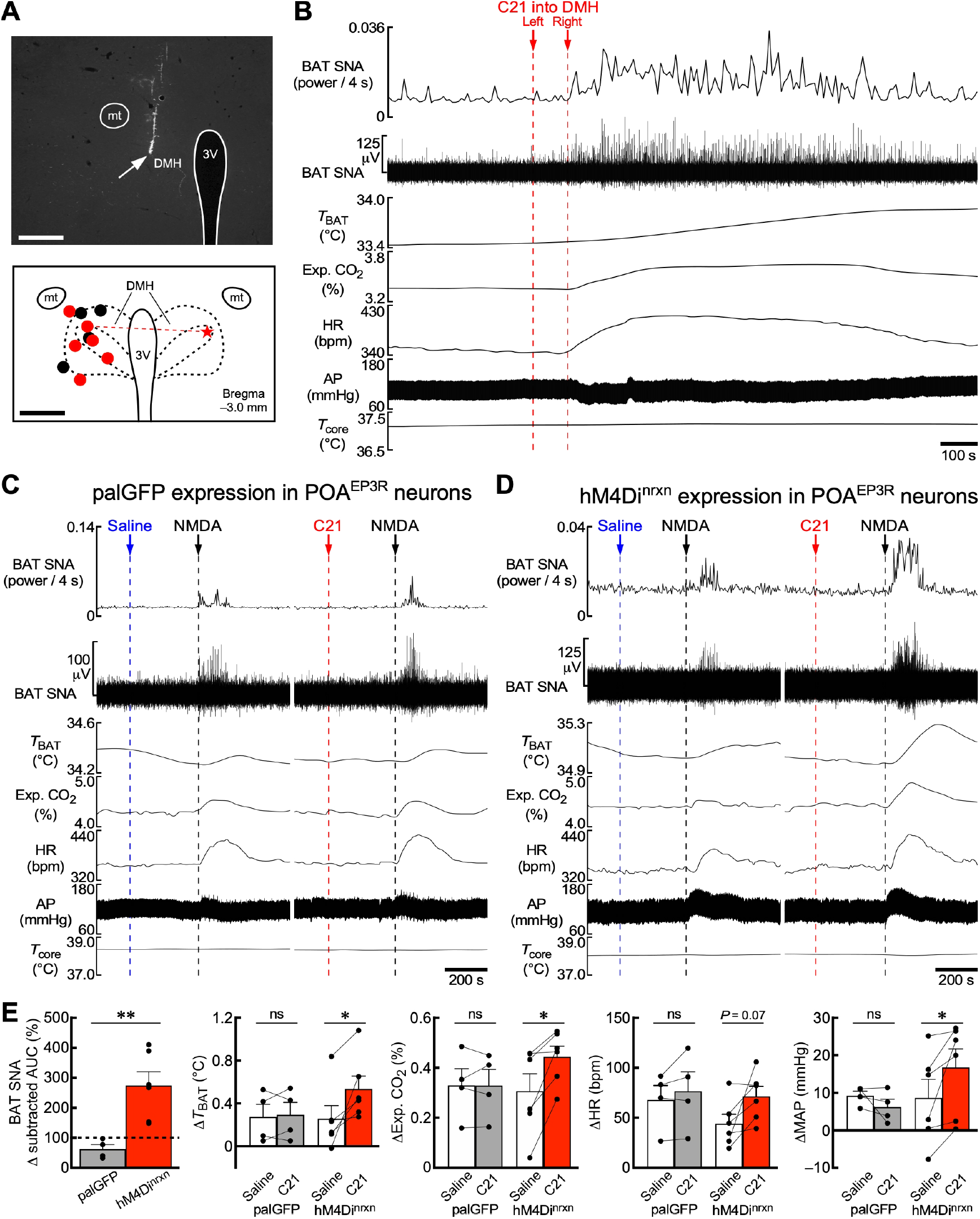
POA^EP3R^→DMH transmission tonically inhibits sympathetic outflow to BAT. (A) A representative view (top) of a nanoinjection site (arrow) in the DMH and a map (bottom) that shows all sites of unilateral injections into the DMH of *Ptger3*-tTA rats with POA^EP3R^ neurons transduced with palGFP (black circles) or mCherry-T2A-hM4Di^nrxn^ (red circles) and bilateral C21 injections shown in (B) (red circle and star tied with a dashed line). Scale bars, 0.5 mm. (B) An example of increases in BAT thermogenesis (BAT SNA and *T*_BAT_), expired (Exp.) CO_2_ and HR evoked by bilateral nanoinjections of C21 into the DMH of a *Ptger3*-tTA rat with POA^EP3R^ neurons transduced with mCherry-T2A-hM4Di^nrxn^. AP, arterial pressure. (C and D) Effects of unilateral nanoinjection of saline or C21 into the DMH of *Ptger3*-tTA rats with POA^EP3R^ neurons transduced with palGFP (C) or mCherry-T2A-hM4Di^nrxn^ (D) on increases in BAT SNA, *T*_BAT_, Exp. CO_2_, HR and AP evoked by NMDA injection at the same DMH site. (E) Group data showing NMDA-induced changes in BAT SNA, *T*_BAT_, Exp. CO_2_, HR and mean arterial pressure (MAP) following saline or C21 injection into the DMH of *Ptger3*-tTA rats with POA^EP3R^ neurons transduced with palGFP (*n* = 4) or mCherry-T2A-hM4Di^nrxn^ (*n* = 6). NMDA-evoked changes in BAT SNA were quantified as the area under the curve (AUC) of the “power/4 s” trace (C and D) above the pre-injection baseline level (subtracted AUC) for 5 min after an NMDA injection. The NMDA-evoked increases in subtracted AUC following C21 injection are expressed as % of NMDA-evoked increases in subtracted AUC following saline injection and compared between the palGFP and mCherry-T2A-hM4Di^nrxn^ groups by an unpaired *t*-test (*t*_8_ = 3.65). NMDA-induced changes in *T*_BAT_, Exp. CO_2_, HR and MAP following saline and C21 injection were compared by paired *t*-tests (*T*_BAT_: palGFP, *t*_3_ = 0.37; hM4Di^nrxn^, *t*_5_ = 3.15; Exp. CO_2_: palGFP, *t*_3_ = 0.02; hM4Di^nrxn^, *t*_5_ = 3.53; HR: palGFP, *t*_3_ = 1.30; hM4Di^nrxn^, *t*_5_ = 2.34; MAP: palGFP, *t*_3_ = 1.47; hM4Di^nrxn^, *t*_5_ = 2.70). ns, not significant; \**P* < 0.05; \*\**P* < 0.01. All values are means ± SEM.

Another potential reason may be that skin warming with a water jacket to maintain *T*_core_ of the anesthetized rats inhibited the POA→DMH glutamatergic excitatory transmission that is required for the excitation of sympathoexcitatory DMH neurons after withdrawal of the GABAergic inhibition from POA^EP3R^ neurons (Tanaka et al., 2011; da Conceição et al., 2020; Piñol et al., 2021). The balance between GABAergic and glutamatergic inputs from the POA to the DMH is considered important for the activity control of the DMH→rMR neurons that drive thermogenic sympathetic outflow (Morrison and Nakamura, 2019; Nakamura et al., 2022). To examine whether POA^EP3R^→DMH neurons contribute to this excitatory/inhibitory (E/I) balance control of sympathetic efferent signaling, we tested the effect of local inhibition of POA^EP3R^→DMH axon terminals on BAT thermogenesis and cardiovascular responses evoked by a glutamatergic excitation in the DMH (Figure 6). A unilateral nanoinjection of NMDA, an ionotropic glutamate receptor agonist, into the DMH consistently elicited increases in BAT sympathetic nerve activity (SNA), *T*_BAT_, expired CO_2_, heart rate (HR) and mean arterial pressure in either *Ptger3*-tTA rats that expressed hM4Di^nrxn^ in POA^EP3R^ neurons or control *Ptger3*-tTA rats that expressed palGFP in POA^EP3R^ neurons (Figures 6C–E). The NMDA-evoked BAT thermogenic and pressor responses were potentiated by a prior nanoinjection of C21 at the same DMH site in hM4Di^nrxn^-expressing rats, but not in control rats (Figures 6C–E). The NMDA-evoked increase in HR tended to be larger following C21 injection than that following saline injection in hM4Di^nrxn^-expressing rats, although statistically insignificant (Figures 6D and 6E). These results support the view that POA^EP3R^→DMH transmission provides tonic inhibitory signals to control sympathoexcitatory DMH neurons and reduction of this tonic inhibition results in an E/I balance shift to enhance the excitability of sympathoexcitatory DMH neurons to drive BAT thermogenesis and cardiovascular responses.

## DISCUSSION

The POA has been considered as a thermoregulatory center as well as a febrile center (Boulant, 2000; Nakamura, 2011). The present study demonstrates that POA^EP3R^ neurons, a target of PGE_2_ for its pyrogenic action, play a pivotal role in the preoptic efferent control of central sympathetic outflow for basal thermoregulation. Heat exposure of rats increased Fos expression in POA^EP3R^ neurons and chemogenetic stimulation of POA^EP3R^ neurons elicited hypothermia at room temperature via a remarkable increase in skin blood flow, an active heat-loss response mediated by sympathoinhibition. On the other hand, chemogenetic inhibition of POA^EP3R^ neurons elicited hyperthermia via an increase in BAT thermogenesis, mimicking fever and cold defense. Therefore, POA^EP3R^ neurons bidirectionally regulate *T*_core_ by controlling both production and dissipation of heat. The present histochemical analyses further showed that POA^EP3R^ neurons provide predominantly GABAergic innervation to sympathoexcitatory DMH neurons whose activation increases *T*_core_ via excitation of sympathetic premotor neurons in the rMR. Our findings provide strong evidence that POA^EP3R^ neurons mediate the tonic inhibitory regulation of sympathoexcitatory efferent pathways, which underlies the fundamental principle of central regulation of body temperature and fever (Figure 7).

**Figure 7.**
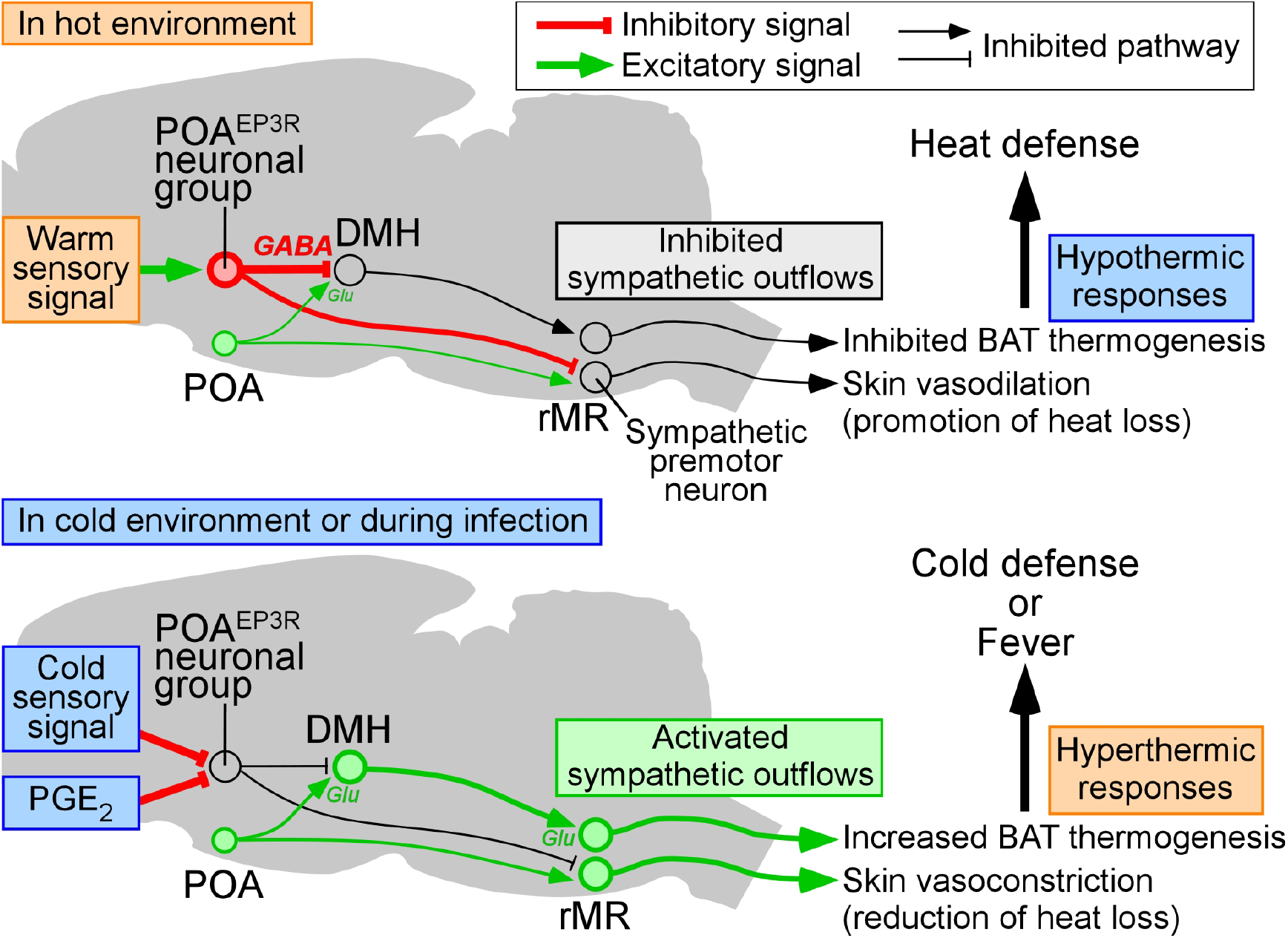
A model of the central circuit mechanism for thermoregulation and fever. This scheme illustrates how tonic inhibitory signaling from POA^EP3R^ neurons bidirectionally controls *T*_core_ for thermal homeostasis and development of fever. In a hot environment (top), cutaneous warm-sensory signals activate POA^EP3R^→DMH projection neurons, which then provide strong GABAergic inputs to sympathoexcitatory DMH neurons to inhibit the sympathetic efferent pathway to BAT and cardiovascular organs. Another separate POA^EP3R^ neuronal subgroup that innervate the rMR may also be activated and inhibit sympathetic outflow to skin vasoconstrictors to elicit skin vasodilation. The inhibitory signaling from POA^EP3R^ neurons dominates over glutamatergic excitatory inputs from the POA (and other as yet unidentified brain regions) to the DMH and rMR. The elicited hypothermic effector responses result in inhibition of heat production and promotion of heat loss for the defense of *T*_core_ in the hot environment. In a cold environment or during infection (bottom), cutaneous cold-sensory signals or PGE_2_ inhibit POA^EP3R^→DMH projection neurons, respectively. Consequently, the GABAergic tonic signaling from POA^EP3R^ neurons is attenuated and thereby, the glutamatergic inputs to the DMH and rMR can drive sympathetic outflows to BAT, skin vasoconstrictors and cardiovascular organs. These sympathetic outputs drive hyperthermic responses, increased BAT thermogenesis and skin vasoconstriction, for the defense of *T*_core_ in the cold environment or for the development of fever.

Earlier studies examined the effects on *T*_core_ of selective stimulation and inhibition of VGLUT2-expressing and VGAT-expressing POA neurons in mice (Song et al., 2016; Yu et al., 2016; Abbott and Saper, 2017; Zhao et al., 2017). However, both glutamatergic and GABAergic populations in the POA are composed of heterogenous subgroups with different projection properties (i.e., interneurons and projection neurons) and different roles in thermoregulation (Morrison and Nakamura, 2019; Nakamura et al., 2022), and therefore, the manipulations of the entire VGLUT2- or VGAT-expressing population of POA neurons have not permitted dissection of the thermoregulatory neural circuit. With efforts to classify POA neuron subgroups, recent studies have identified various genetic markers: pituitary adenylate cyclase-activating polypeptide (PACAP), brain-derived neurotrophic factor (BDNF), leptin receptor (LepRb), galanin, neuronal nitric oxide synthase (NOS1), pyroglutamylated RFamide peptide (QRFP), estrogen receptor α (ERα) and neuropsin (Opn5) (Tan et al., 2016; Yu et al., 2016; Harding et al., 2018; Kroeger et al., 2018; Ma et al., 2019; Hrvatin et al., 2020; Takahashi et al., 2020; Zhang et al., 2020a, 2020b). Stimulation of any of these POA neuron subgroups results in hypothermia at room temperature and some of them induces torpor-like severe hypothermia with *T*_core_ reduced to room temperature (Hrvatin et al., 2020; Takahashi et al., 2020; Zhang et al., 2020a, 2020b). However, none of them elicits obvious hyperthermia when inhibited. Therefore, these subgroups likely mediate heat defense and some of them may drive torpor when mice are hungry in cold environments, but they seem unlikely to function for cold defense or fever. Another group of POA neurons expressing bombesin-like receptor 3 (BRS3) is activated by cooling of mice and drive cold-defensive responses, but do not mediate LPS-induced fever (Piñol et al., 2021). Chemogenetic inhibition of BRS3-expressing POA neurons elicits only a modest decrease (0.5°C) in *T*_core_ (Piñol et al., 2021). In contrast, the present study showed that POA^EP3R^ neurons, which are activated by heat exposure and inhibited by PGE_2_, bidirectionally control *T*_core_. The chemogenetic stimulation of POA^EP3R^ neurons reduced *T*_core_ by 1.9°C, which was less severe than the torpor-like hypothermia induced by the other POA neuron subgroups. Therefore, POA^EP3R^ neurons constitute a unique and pivotal group of thermoregulatory POA neurons that mediates not only fever development, but also thermal homeostasis in both hot and cold environments.

Because POA^EP3R^ neurons exhibited a low level of Fos expression during exposure to room temperature, we could not detect cold exposure-induced inhibition of these neurons by monitoring Fos, which is an immediate-early transcription factor expressed in response to intense neuronal excitation involving an elevation of intracellular Ca^2+^ (Sheng et al., 1990). However, chemogenetic inhibition of POA^EP3R^ neurons at room temperature elicited BAT thermogenesis and hyperthermia. This result indicates that POA^EP3R^ neurons under thermoneutral conditions exhibit a certain level of tonic firing that does not involve Ca^2+^ influx leading to Fos expression, and inhibition of this tonic activity, probably by a cold-sensory input from the skin through the lateral parabrachial nucleus (Nakamura and Morrison, 2008), elicits cold-defensive thermoregulatory responses including BAT thermogenesis. Future electrophysiological recordings from POA^EP3R^ neurons will allow us to directly measure their activity responses to cutaneous thermosensory inputs and, also interestingly, to determine whether these neurons have intrinsic thermosensitivity as warm-sensitive neurons that sense local brain tissue temperature (Nakayama et al., 1961).

The increased activity of POA^EP3R^ neurons by heat exposure was reduced by PGE_2_. Although the EP3R is mostly coupled to Gi in culture cells (Narumiya et al., 1999), it has been unknown how PGE_2_ alters POA neuronal activities through the EP3R to trigger fever. Our results demonstrate that PGE_2_ inhibits POA^EP3R^ neuronal activity to elevate *T*_core_, and a correlation analysis between Fos expression and *T*_core_ predicted that suppression of all POA^EP3R^ neurons by PGE_2_ would increase *T*_core_ up to 42.1°C, close to the typical upper limit of fever (Mackowiak and Boulant, 1996). The view that the Gi-mediated reduction of the intracellular cAMP levels in POA^EP3R^ neurons inhibits their neural activity and thereby mediates the febrile action of PGE_2_ is consistent with previous findings that intracerebroventricular injection of PGE_2_ reduces cAMP levels in the POA and blockade of cAMP degradation in the POA impairs PGE_2_- induced fever (Steiner et al., 2002).

An intriguing finding in this study is the discrepant expressions of the transmitter markers in the cell bodies and axon terminals of POA^EP3R^ neurons. The glutamatergic neuronal marker, VGLUT2 mRNA, was more prevalently expressed in their cell bodies than the GABAergic marker, GAD67 mRNA, consistent with a recent finding that ablation of the *Ptger3* gene in Cre-expressing cells in VGLUT2-Cre mice results in blunted febrile response to LPS (Machado et al., 2020). However, our anterograde tracing from POA^EP3R^ neurons combined with immunohistochemistry for VGLUT2 and VGAT proteins revealed that POA^EP3R^ neurons predominantly release GABA, rather than glutamate, from their axon terminals in the DMH. The DMH contains sympathoexcitatory neurons that innervate the rMR to drive cold-defensive, febrile and stress responses (Zaretskaia et al., 2003; Madden and Morrison, 2004; Nakamura et al., 2005; Nakamura and Morrison, 2011; Kataoka et al., 2014, 2020), and we found that cell bodies of rMR-projecting DMH neurons were more often in close contact with GABAergic axons of POA^EP3R^ neurons than glutamatergic axons. Our immunohistochemical procedure was validated by the results that VGAT-immunoreactive puncta were co-labeled with other antibodies raised against different epitopes on the VGAT protein and that our VGLUT2 immunohistochemistry successfully visualized glutamatergic terminals in palGFP-labeled thalamocortical axons. Furthermore, many VGAT-immunoreactive puncta of POA^EP3R^ neuronal axons in the DMH were accompanied by GAD67 immunoreactivity, indicating that these axon terminals can both produce and release GABA. Combined with our physiological data, these observations strongly support the central thermoregulatory and febrile mechanisms, in which POA^EP3R^ neurons directly innervate DMH→rMR sympathoexcitatory neurons via tonic GABAergic inputs, and attenuation of the tonic GABAergic signals by cutaneous cold-sensory inputs to or by a PGE_2_ action on POA^EP3R^ neurons disinhibit the sympathoexcitatory neurons to drive thermogenic and cardiovascular responses (Figure 7).

Our observation that EP3Rs are expressed by both VGLUT2 mRNA-expressing and GAD67 mRNA-expressing POA neuronal groups is consistent with the recent single cell/nucleus RNA sequencing data showing *Ptger3* mRNA expression in “excitatory” and “inhibitory” POA cell clusters (Moffitt et al., 2018; Upton et al., 2021). The abundant GABAergic terminals of POA^EP3R^→DMH axons might be provided exclusively by the GAD67 mRNA-expressing population. However, DMH-projecting POA^EP3R^ neurons predominantly express VGLUT2 mRNA (Figure S3G). Therefore, VGLUT2 mRNA-expressing POA^EP3R^ neurons also likely produce the proteins required to be GABAergic, such as VGAT and GAD67. It is important to note that mRNA transcript levels by themselves are not sufficient to predict their protein product levels in many types of cells (Liu et al., 2016), and therefore, in VGLUT2 mRNA-expressing POA^EP3R^ neurons, sufficient amounts of such GABAergic presynaptic proteins could be translated from low levels of mRNA transcripts. Furthermore, these mRNAs may be targeted to axon terminals for local translation into those GABAergic presynaptic proteins (Jung et al., 2012; Holt et al., 2019) and such axonal mRNAs would be difficult to be detected by *in situ* hybridization or single cell/nucleus RNA sequencing. The present results provide a potential caveat that VGLUT2 gene transcripts in the cell bodies, particularly in POA neurons, are not a reliable marker for glutamatergic neurons.

Another interesting finding is the increase in VGAT-immunoreactive puncta of POA^EP3R^→DMH axons following chronic heat exposure. This observation leads us to hypothesize a heat adaptation mechanism, in which POA^EP3R^ neurons during chronic heat exposure form more GABAergic inhibitory synapses onto sympathoexcitatory DMH neurons to increase the ability to tolerate heat stress. The concomitant reduction of POA^EP3R^→DMH axon swellings lacking VGAT and VGLUT2 suggests that these ‘empty’ axons serve as a reserve, which is to be installed with GABAergic presynaptic machinery when necessary to tolerate heat. The local translation in axon terminals may be involved in the installation with GABAergic presynaptic proteins, and how chronic heat exposure triggers this adaptation mechanism is an interesting question for future research.

In the present study, local chemogenetic inhibition of transmitter release from POA^EP3R^→DMH axon terminals potentiated BAT thermogenic, metabolic and pressor responses evoked by glutamatergic stimulation of DMH neurons. In some rats, bilateral chemogenetic inhibition of POA^EP3R^→DMH axon terminals by itself elicited robust activation of BAT thermogenesis and tachycardia. Together with the histochemical observations, these physiological results indicate that the POA^EP3R^→DMH tonic GABAergic signal is a critical determinant of the E/I balance that regulates the excitability of sympathoexcitatory DMH neurons whose activation leads to elevation of *T*_core_. The hyperthermia evoked by chemogenetic inhibition of POA^EP3R^ neurons in awake rats (Figure 5D) suggests that sympathoexcitatory DMH neurons receive a certain level of glutamatergic inputs in awake conditions under thermoneutral environments, but cannot actively generate action potentials until concurrent tonic GABAergic inputs from POA^EP3R^ neurons are attenuated by cold-sensory inputs or PGE_2_ (Figure 7). This mechanism is consistent with the view that the POA^EP3R^→DMH GABAergic signaling, whose tone is altered by cutaneous and central thermosensory signals, makes a major contribution to setting the control range of *T*_core_. The glutamatergic inputs from the POA to sympathoexcitatory DMH neurons may also be regulated by thermosensory signals (activated by cold) (Tanaka et al., 2011; da Conceição et al., 2020; Piñol et al., 2021). However, sympathoexcitatory DMH neurons also likely receive temperature-independent, intense excitatory signals from other brain sites perhaps located caudal to the POA, because a complete disruption of fibers descending from the POA elicits robust BAT thermogenesis and cutaneous vasoconstrictor activity leading to hyperthermia (Chen et al., 1998; Rathner et al., 2008). The present physiological results indicate that the POA^EP3R^→DMH pathway controls BAT thermogenesis and accompanying cardiovascular responses. This conclusion is supported by the finding that inactivation of DMH neurons abolishes BAT thermogenesis and cardiovascular responses for fever and cold defense (Madden and Morrison, 2004; Nakamura et al., 2005; Nakamura and Morrison, 2007). On the other hand, the DMH does not mediate cutaneous vasoconstriction for fever or cold defense (Rathner et al., 2008), although it does mediate psychological stress-induced cutaneous vasoconstriction (Kataoka et al., 2020). Another parallel pathway from POA^EP3R^ neurons to the rMR likely controls cutaneous vasomotion for basal thermal homeostasis and fever (Nakamura et al., 2002; Tanaka et al., 2011) (Figure 7). The rMR-projecting subgroup of POA^EP3R^ neurons is distinct from that projecting to the DMH (Nakamura et al., 2009). Although the present anterograde tracing visualized the distribution of POA^EP3R^ neuronal axons in the rMR, their transmitter properties remain to be investigated. Further studies are required to determine how these two descending pathways from POA^EP3R^ neurons cooperatively regulate the effectors for thermoregulation and fever development.

## EXPERIMENTAL PROCEDURES

### Generation of *Ptger3*-tTA BAC transgenic rats

We modified a BAC clone (CH230-35G1) from a female BN/SsNHsd/MCW rat genomic BAC library (BACPAC Resources) that consisted of pTARBAC2.1 vector and a genomic DNA fragment (∼ 243 kb) containing the *Ptger3* gene. To replace the loxP site in the BAC vector with a homing endonuclease (I-*Sce*I) recognition site, we first generated pTARBAC1H vector from pTARBAC1 vector (BACPAC Resources) by replacing the loxP site with I-*Sce*I site and by deleting the loxP511 site. The pTARBAC2.1 backbone of the BAC clone was then replaced with the pTARBAC1H backbone by Red/ET recombination (Counter Selection BAC modification kit; Genebridges, Heidelberg, Germany) using an appropriate rpsL/neo cassette according to manufacturer’s manual. The recombined BAC DNA was further subjected to another Red/ET recombination to insert a DNA cassette (989 bp) consisting of tTA Advanced (tTAad), an improved version of tetracycline-controlled transactivator, and bovine growth hormone polyadenylation sequence (BGH) at immediately 3’ to the start ATG codon of the *Ptger3* gene (Figure S4A).

The recombined *Ptger3*-tTA-BGH BAC was purified with a Nucleobond BAC100 kit (Macherey-Nagel, Düren, Germany) and the DNA was digested for 1 h with I-*Sce*I (New England BioLabs, Beverly, MA) at 37°C. The linearized BAC DNA was then injected into pronuclei of Sprague-Dawley rat zygotes (Japan SLC, Shizuoka, Japan). The zygotes were transferred into pseudo-pregnant female rats, and 63 pups were obtained. Genomic DNAs of 61 weaned pups (2 pups died before weaning) were collected from their tails with a DNeasy Tissue kit (Qiagen, Valencia, CA). Six *Ptger3*-tTA BAC transgenic founder rats were identified by PCR for the presence of the tTAad-BGH insert, 5’-end (T7 arm) and 3’-end (Sp6 arm) of the linearized BAC DNA by PCR (Figure S4B). To further evaluate the selectivity of reporter gene expression in these founder lines, we injected AAV-TRE-palGFP into the POA and performed double immunostaining for GFP and EP3R (see below). One of the founder lines, which exhibited the highest specificity of palGFP expression in POA^EP3R^ neurons (line ID: 101001-09-5; Figures 4A and 4B), was chosen and used in the present study. Based on the inheritance pattern of the transgene, in this founder line, the BAC transgene was found to be located in the X chromosome, but the transgenic rats were fertile with no obvious abnormalities. The transgenic line was maintained in heterozygotes on a Sprague-Dawley genetic background, and transmission of the transgene to offsprings was monitored by PCR.

### Animals

Male Sprague-Dawley rats (200–500 g; Japan SLC) and male *Ptger3*-tTA rats (200– 500 g) were used. They were housed two or three to a cage with *ad libitum* access to food and water in a room air-conditioned at 25 ± 2°C with a standard 12 h light/dark cycle (lights on 07.00–19.00 h) before used for surgery or experiments. All procedures conform to the guidelines of animal care by the Division of Experimental Animals, Nagoya University Graduate School of Medicine and were approved by the Nagoya University Animal Experiment Committee.

### AAV vectors

Production of AAV vectors followed our established procedure (Kataoka et al., 2020; Takahashi et al., 2021). For production of pAAV2-TRE-palGFP, a palGFP DNA cassette was amplified with a PCR method using pAAV-CMV-palGFP (Kataoka et al., 2014) as a template together with primers containing 5’-*BamH*I site and 3’-*Mlu*I site. After digested with these restriction enzymes, this cassette was inserted into the multi-cloning site of pENTR1A-TRE (Hioki et al., 2009) between the *BamH*I/*Mlu*I sites. Similarly, for production of pAAV2-TRE-mCherry-T2A-hM4Di^nrxn^, an mCherry-T2A-hM4Di^nrxn^ cassette was PCR-amplified using CAG::mCherry-2a-hM4Dnrxn (Addgene #52523, donated by Dr. Scott Sternson) as a template with primers containing 5’-*BamH*I site and 3’-*Mlu*I site and then inserted into pENTR1A-TRE. For production of pAAV2-TRE-hM3Dq-mCherry, an hM3Dq-mCherry cassette was PCR-amplified using pAAV-hSyn-DIO-hM3Dq-mCherry (Addgene #44361, donated by Dr. Bryan Roth) as a template with primers containing 5’-*Mlu*I site and 3’-*EcoR*I site and then inserted into pENTR1A-TRE between the *Mlu*I/*EcoR*I sites. The produced entry vectors, pENTR1A-TRE-palGFP, pENTR1A-TRE-mCherry-T2A-hM4Di^nrxn^ and pENTR1A-TRE-hM3Dq-mCherry, were subjected to homologous recombination using LR clonase II with the destination vector, pAAV2-DEST(r) (Sohn et al., 2017) to produce pAAV2-TRE-palGFP, pAAV2-TRE-mCherry-T2A-hM4Di^nrxn^ and pAAV2-TRE-hM3Dq-mCherry. The pAAV2 vectors were used for production and purification of AAV2/1-TRE-palGFP, AAV2/1-TRE-mCherry-T2A-hM4Di^nrxn^ and AAV2/1-TRE-hM3Dq-mCherry according to our methods (Kataoka et al., 2020; Takahashi et al., 2021). The final titrations were 1.2 × 10^13^ GC/ml (AAV-TRE-palGFP), 1.1 × 10^13^ GC/ml (AAV-TRE-mCherry-T2A-hM4Di^nrxn^) and 1.3 × 10^13^ GC/ml (AAV-TRE-hM3Dq-mCherry). AAV-TRE-hM3Dq-mCherry was diluted 50-fold with 0.9% saline before intracranial injection.

### Stereotaxic injection

Rats were anesthetized with a combination anesthetic (0.15 mg/kg medetomidine hydrochloride, 2.0 mg/kg midazolam, 2.5 mg/kg of butorphanol tartrate; i.p. or s.c.) following gas anesthesia with 3% isoflurane and were positioned in a stereotaxic apparatus. A glass micropipette filled with a solution containing AAV, CTb conjugated with Alexa Fluor 488 (Alexa488) or Alexa647 (1 mg/ml, C22841 and C34778, Thermo Fisher) or FG (4% solution dissolved in 0.9% saline; Fluorochrome, Denver, CO) was perpendicularly inserted into the POA (AAV), anterior midline thalamus (AAV), DMH (CTb or FG) or rMR (FG). The solution was pressure-ejected by using a Picospritzer III (Parker, Hollis, NH), and the micropipette was remained for 5 min after injection before it was withdrawn. AAV injections into the POA targeted at 3 sites (150 nl/site): MnPO (0.3 mm rostral to bregma, on the midline, 7.2 mm ventral to the brain surface) and MPA (0.1 mm rostral to bregma, 0.6–0.7 mm left and right to the midline, 7.8 mm ventral to the brain surface). AAV injection into the anterior midline thalamus (200 nl) was made at the coordinates: 2.0 mm caudal to bregma, on the midline and 5.0–5.5 mm ventral to the brain surface. CTb injection (250 nl) was made unilaterally into the DMH (3.1 mm caudal to bregma, 0.4–0.8 mm lateral to the midline and 8.5 mm ventral to the brain surface). FG injections into the DMH were made at the same coordinates but bilaterally (50 nl/site). FG injection into the rMR (80 nl) was made at the coordinates: 2.8 mm caudal to the interaural line, on the midline and 9.6 mm ventral to the brain surface. After the injection, all incisions were sutured and disinfected with iodine, and ampicillin sodium (0.2 ml, 125 mg/ml) and atipamezole hydrochloride (250 μg/kg) solutions were injected into femoral muscles. The rats were housed 7–10 days under regular health check until subsequent procedure or experiments.

### Cold and heat exposure

Cold and heat exposure of rats was performed in a climate chamber (Nakamura et al., 2004) with the standard light/dark control as described above. Rats were individually placed in plastic cages with wire mesh lids with *ad libitum* food and water. For acute exposure experiments, the cages were placed in the climate chamber air-conditioned at 24°C overnight to acclimatize the rats to the chamber, and then, they were exposed to 4°C, 24°C or 36°C for 2 hrs (approximately between 09.00 and 12.00 h). For chronic exposure, rats were exposed to 10°C, 25°C or 34°C for 16 days (10°C) or 14 days (25°C and 34°C), except short periods of time for regular health check and cage maintenance. Immediately after the exposure, the rats were anesthetized and transcardially fixed for histological analyses as described below.

### Immunohistochemistry

Immunohistochemical procedures followed our previous studies (Nakamura et al., 2000, 2004). Rats were anesthetized and transcardially perfused with saline and then with 4% formaldehyde in 0.1 M phosphate buffer (pH 7.4). The brain was removed, postfixed in the fixative at 4°C for 2–3 hrs, and then cryoprotected in a 30% sucrose solution longer than overnight. The tissue was cut into 30-µm-thick frontal sections on a freezing microtome. The primary antibodies used are anti-EP3R rabbit antibody (1 μg/ml; Nakamura et al., 1999, 2000), anti-Fos goat antibody (1:1000; sc-52G, Santa Cruz Biotechnology), anti-CTb goat serum (1:5,000; #703, List Biological Laboratories), anti-GFP mouse antibody (1:200; A11120, Thermo Fisher), anti-GFP rabbit antibody (0.5 μg/ml; Tamamaki et al., 2000), anti-VGAT guinea pig serum (1:1,000; 131004, Synaptic Systems), anti-VGLUT2 rabbit antibody (0.5 μg/ml; Hioki et al., 2003), anti-GAD67 mouse antibody (1:300; MAB5406, Sigma-Aldrich), anti-synaptophysin mouse antibody (1:1,000; S5768, Sigma-Aldrich) and anti-mRFP guinea pig antibody (1 μg/ml; Hioki et al., 2010). The anti-mRFP antibody showed cross-reactivity to mCherry. VGAT immunohistochemistry using the anti-VGAT guinea pig serum (epitope: amino acids 2–115 on rat VGAT) was validated by co-staining with anti-VGAT mouse antibody (1:200; 131011, Synaptic Systems; epitope: amino acids 75–87 on rat VGAT) and anti-VGAT rabbit antibody (1:200; VGAT11-A, Alpha Diagnostics; epitope: 17-amino acids of the C-terminus of rat VGAT).

For double immunofluoscence staining for EP3R and Fos, sections were incubated with anti-EP3R rabbit antibody and anti-Fos goat antibody overnight at 4°C and then with Alexa594-conjugated donkey antibody to goat IgG (10 μg/ml; A11058, Thermo Fisher) and biotinylated donkey antibody to rabbit IgG (1:100; AP182B, Merck Millipore) for 1 hr at room temperature. After rinsed, these sections were further incubated with avidin-biotinylated peroxidase complex (ABC; 1:50; PK-6100, Vector Laboratories) for 1 hr and then, EP3R immunoreactivity was visualized by reaction with FITC-conjugated tyramide (1:50; SAT701001KT; Tyramide Signal Amplification FITC Systems, PerkinElmer) for 3 min. For double immunofluoscence staining for EP3R and mCherry, immunoreactivity for mCherry was detected by anti-mRFP guinea pig antibody and Alexa594-conjugated goat antibody to guinea pig IgG (10 μg/ml; A11076, Thermo Fisher).

For triple immunolabeling in palGFP-labeled axons, brain sections were incubated overnight with a mixture of anti-GFP mouse antibody, anti-VGAT guinea pig serum and anti-VGLUT2 rabbit antibody, a mixture of anti-GFP rabbit antibody, anti-VGAT guinea pig serum and anti-GAD67 mouse antibody, a mixture of anti-GFP rabbit antibody, anti-VGAT guinea pig serum and anti-synaptophysin mouse antibody, or a mixture of anti-VGAT guinea pig serum, anti-VGAT mouse antibody and anti-VGAT rabbit antibody. The sections were then incubated for 2 hrs with an appropriate combination of the secondary antibodies (10 μg/ml for each): Alexa488-conjugated goat antibody to mouse IgG (A11029, Thermo Fisher), Alexa488-conjugated goat antibody to rabbit IgG (A11034, Thermo Fisher), Alexa568-conjugated goat antibody to guinea pig IgG (A11075, Thermo Fisher), Alexa647-conjugated goat antibody to rabbit IgG (A21245, Thermo Fisher), and Alexa647-conjugated goat antibody to mouse IgG (A21236, Thermo Fisher). Immunoreactivity of the anti-VGAT rabbit antibody was detected by incubation with biotinylated donkey antibody to rabbit IgG followed by incubation with streptavidin-conjugated Alexa647 (1:400; S21374, Thermo Fisher).

### Fluorescence *in situ* hybridization

Rats were anesthetized and transcardially perfused with 4% formaldehyde in 0.1 M phosphate buffer (pH 7.4). The brains were postfixed in the fixative at 4°C for 3 days, and then cryoprotected in a 30% sucrose solution for at least 2 days. The tissues were cut into 35-µm-thick frontal sections on a freezing microtome in an RNase-free environment. These sections were blocked in a hybridization buffer [4.95 x saline sodium citrate (SSC), 1.98% Blocking Reagent (11096176001, Roche), 49.5% formamide, 0.099% N-lauroylsarcosine (NLS), 0.099% sodium dodecyl sulfate] at 60°C for 1 hr and then incubated with 1 μg/ml of digoxigenin-labeled, rat GAD67 or mouse VGLUT2 antisense RNA probe (Nakamura et al., 2002, 2007) in this buffer at 60°C for 20–22 hrs. The sections were washed with 2 x SSC containing 50% formamide and 0.1% NLS at 60°C for 20 min twice, and then further washed with the following solutions at 37°C for 20 min twice for each solution: 1) 2 x SSC containing 0.1% NLS and 2) 0.2 x SSC containing 0.1% NLS. After rinsed with TS7.5 solution [0.1 M Tris-HCl, 0.15 M NaCl (pH7.5)] at room temperature for 5 min, the sections were blocked in TS7.5 containing 1% Blocking Reagent for 1 hr and then incubated with alkaline phosphatase-conjugated anti-digoxigenin sheep antibody (1:1000; 11093274910, Roche) and anti-EP3R rabbit antibody in this blocking solution overnight at 4°C. In case of detecting Fos, anti-Fos goat antibody was added to this antibody cocktail. After rinsed with TS7.5 containing 0.05% Tween 20 (TNT) for 5– 10 min twice, these sections were incubated with biotinylated donkey antibody to rabbit IgG in TS7.5 containing 1% Blocking Reagent for 1 hr at room temperature. In case of detecting Fos immunoreactivity, Alexa647-conjugated donkey antibody to goat IgG (10 μg/ml; A21447, Thermo Fisher) was added to this solution. After rinsed with TNT for 5 min twice, the sections were incubated with TS7.5 containing ABC (1:50) for 1 hr. After washed with TNT for 10 min twice and then with TS8.0 solution [0.1 M Tris-HCl, 0.1 M NaCl, 10 mM MgCl_2_ (pH8.0)] for 10 min twice, the sections were incubated with an HNPP/Fast Red TR solution (11758888001, Roche) at room temperature for 2.5 hr. After rinsed with TS7.5, these sections were further incubated with FITC-conjugated tyramide as described above. These sections were rinsed with TS7.5 and then soaked in ice-cold TS8.0 to be mounted onto glass slides. The slides were dried in the dark and then, briefly soaked into TS8.0 to be coverslipped with CC/Mount (Diagnostic BioSystems). In our preliminary experiments, we confirmed that the mouse VGLUT2 antisense probe showed hybridization signals in the rat brain consistent with reported VGLUT2 mRNA distribution (Nakamura et al., 2007).

For fluorescence *in situ* hybridization combined with EP3R and CTb immunohistochemistry, hybridization signals were detected using peroxidase instead of alkaline phosphatase. Sections after the post-hybridization washing and blocking steps were incubated with peroxidase-conjugated anti-digoxigenin sheep antibody (1:1000; 11207733910, Roche) in TS 7.5 containing 1% Blocking Reagent at 4°C for overnight. After rinsed with TS7.5, the sections were incubated with Cy3-conjugated tyramide (1:50) in Amplification Diluent (SAT704A001KT, PerkinElmer) supplemented with 0.00014% of H_2_O_2_ and 4% of polyvinyl alcohol (Mw 31,000– 50,000; 363138, Sigma-Aldrich) for 6 hrs at room temperature. After washed with TNT for 30 min, the sections were incubated in phosphate-buffered saline (PBS) containing 3% of H_2_O_2_ for 10 min at room temperature to block the peroxidase activity. The sections were thoroughly washed with TNT and with TS 7.5, and then incubated with anti-EP3R rabbit antibody and anti-CTb goat serum in TS 7.5 containing 1% of Blocking Reagent at 4°C for overnight. After rinsed with TNT, the sections were further incubated with Alexa647-conjugated donkey antibody to goat IgG (for rats injected with Alexa647-conjugated CTb) or Alexa488-conjugated donkey antibody to goat IgG (10 μg/ml; A11055, Thermo Fisher) (for rats injected with Alexa488-conjugated CTb) in TS 7.5 containing 1% of Blocking Reagent at room temperature for 1 hr. The sections were blocked with 10% normal goat serum in TS7.5 for 30 min and then incubated with biotinylated donkey antibody to rabbit IgG in TS7.5 containing 1% Blocking Reagent and 10% normal goat serum for 1 hr at room temperature. The sections were further incubated with ABC in TS7.5 for 1 hr and then, EP3R immunoreactivity was visualized by reaction with FITC-conjugated tyramide as described above (for rats injected with Alexa647-conjugated CTb) or with Alexa647-conjugated tyramide (1:50–1:100; B40958, Thermo Fisher) in PBS containing 0.0002– 0.0015% of H_2_O_2_ for 5 min (for rats injected with Alexa488-conjugated CTb). The sections were mounted on glass slides and dried in the dark at 4°C overnight.

### Microscopy and histological quantification

Stained sections mounted on glass slides were observed under an epifluorescence microscope (Eclipse 80i, Nikon) or a confocal laser-scanning microscope (TCS SP8, Leica). FG was detected with the fluorescence of the tracer. Confocal images were acquired using the z-stacking function at an interval of 2.5 μm for cell body images in the POA or 1.8 μm for axon images in the DMH and tiled with a LAS X software (Leica).

Labeled cell bodies in EP3R-immunoreactive areas (Figure S1A) were mapped in every sixth section through the POA (representative examples are shown in Figures 1C, S1B, S1D, S2, S3A, S3C and S3D) and manually counted. The quantification was made bilaterally, except for CTb-labeled cell bodies (Figures 3H and S3G), which were counted in the MnPO and the MPA unilateral to the CTb injection. For quantification of presynaptic structures of DMH-projecting axons of POA^EP3R^ neurons, confocal z-stack images covering the entire DMH were acquired bilaterally at two different rostrocaudal levels (5 sections apart) including bregma –3.1 mm (Figure S7D). For quantification of thalamocortical axon terminals, confocal z-stack images of the area 2 of the cingulate cortex were acquired unilaterally from one section. After tiling these z-stack images, we counted puncta immunoreactive for the presynaptic vesicular transporter proteins, VGAT and VGLUT2, in palGFP-labeled axons, which were regarded as presynaptic sites that release GABA and glutamate, respectively. To quantify palGFP-labeled axon swellings that contained neither VGAT-nor VGLUT2-immunoreactive profile (‘empty’ swellings), we counted axon swellings that were at least 1.5 times thicker than the adjoining axon shaft. Data from animals where injection was not centered in the target brain region were excluded.

### Telemetry monitoring and drug injection in free-moving rats

Surgical procedures followed our previous study (Kataoka et al., 2014). *Ptger3*-tTA rats were anesthetized and positioned in a stereotaxic apparatus. After AAV was injected into the POA as described above, a sterile stainless guide cannula (ID = 0.39 mm, OD = 0.71 mm; C313G; Plastic One, Roanoke, VA) was perpendicularly inserted to target the right lateral ventricle (coordinates: 0.8 mm caudal to bregma, 1.8 mm lateral to the midline, 2.5 mm ventral to the brain surface). The inserted guide cannula was anchored with dental cement to stainless steel screws attached to the skull. A dummy cannula cut to the exact length of the guide cannula was inserted into the guide cannula to avoid clogging. The incisions in the skin were closed with suture and the wounds were treated with iodine.

One week after the cannulation, the rats were re-anesthetized and implanted with a battery-operated telemetric transmitter that projected two cables of external thermistor probes (F40-TT, Data Science International, St Paul, MN). The transmitter body and one of the thermistor probes to measure *T*_core_ were placed in the abdominal cavity and sutured to the abdominal wall. The other probe was brought to the back through a tunnel under the skin, inserted into the interscapular BAT pad to measure *T*_BAT_, and tied to the fat tissue with suture. We were careful that the whole part of the probe head for *T*_BAT_ was completely buried into the adipose tissue, but was not in contact with the muscle underneath the BAT pad. All the incisions in the abdominal cavity and skin were closed with suture and the wounds were treated with iodine. After every surgery, the rats were intramuscularly administered with ampicillin sodium and atipamezole hydrochloride. They were housed individually for > 1 week to recover from the surgery under regular health check. During the recovery period, the rats were habituated to the experimenters and injection procedures once every day.

An internal cannula for injection with the thickness to fit the guide cannula was cut to be long enough to allow the injector tip to protrude 1.0 mm below the tip of the guide cannula. The other end of the internal cannula was connected to a polyethylene tubing whose other end was connected to a Hamilton syringe. The inside of the cannula, tubing and syringe was filled with pyrogen-free 0.9% saline (Otsuka, Tokyo, Japan) or C21 (11-(1-piperazinyl)-5*H*-dibenzo[*b,e*][1,4]diazepine dihydrochloride; 200 µg/ml dissolved in saline; SML2392, Sigma-Aldrich). C21 is a potent and selective agonist of hM3Dq and hM4Di and an alternative to the conventional actuator, clozapine-*N*-oxide, as C21 has no reported back-metabolism to clozapine *in vivo* (Thompson et al., 2018). The dose of C21 we used was in the range of effective *in vivo* doses determined in mice (Thompson et al., 2018). The rats were lightly anesthetized with isoflurane, the dummy cannula was gently removed, and the internal cannula was inserted into the guide cannula. The solution (5 µl) was ejected into the lateral ventricle using the syringe at 10.00 h. The rats recovered from anesthesia soon after injection, and *T*_core_, *T*_BAT_ and activity were monitored with a telemetry system (Data Science International) in a room air conditioned at 25 ± 1°C. Each rat received saline and C21 injections at an interval of > 1 week.

In some rats, tail skin temperature was measured using a thermal imaging camera (FLIR C2, FLIR). Snapshot thermographic images were taken right before isoflurane anesthesia and 1 hr after saline or C21 injection. FLIR tools plus software was used to acquire skin temperature at 2–3 cm from the base of the tail.

In experiments involving PGE_2_ injection (Figure 2A), Sprague-Dawley rats were cannulated in the right lateral ventricle as above and implanted with a telemetric transmitter for *T*_core_ recording in the abdominal cavity (TA-F40, Data Science International, St Paul, MN). After a recovery period of > 1 week, telemetry monitoring of *T*_core_ was initiated in a room air conditioned at 25°C. The rats were lightly anesthetized with isoflurane and 5 µl of pyrogen-free 0.9% saline or PGE_2_ (1 mg/ml dissolved in saline; P5640, Sigma-Aldrich) was injected into the lateral ventricle through the cannula as mentioned above. The rats quickly recovered from anesthesia, and 10 min after the injection, they were placed in a climate chamber air-conditioned at 36°C. After monitoring of their *T*_core_ in the chamber for 75 min, the rats were immediately anesthetized and transcardially perfused as above.

### *In vivo* electrophysiology in anesthetized rats

*In vivo* physiological recordings followed our established procedure (Nakamura and Morrison, 2007, 2011). *Ptger3*-tTA rats that had recovered from the surgery for AAV injections into the POA were anesthetized intravenously with urethane (0.8 g/kg) and α-chloralose (80 mg/kg) after cannulation of a femoral artery, a femoral vein, and the trachea under anesthesia with 3% isoflurane in 100% O_2_. They were then placed in a stereotaxic apparatus, and the arterial cannula was attached to a pressure transducer to record arterial pressure and HR. *T*_core_ was monitored from the rectum and maintained at 36.0–38.0°C by perfusing a plastic water jacket, which was wrapped around the shaved trunk, with warm or cold water. The animal was then artificially ventilated with 100% O_2_ through the tracheal cannula and paralyzed with D-tubocurarine (0.6 mg i.v. initial dose, supplemented with 0.3 mg/hr) to stabilize BAT nerve recording by preventing respiration-related movements. Every time the effect of tubocurarine waned, the depth of anesthesia was re-assessed prior to supplementation of tubocurarine and the anesthetic was supplemented as necessary. Mixed expired CO_2_ was monitored through the tracheal cannula using a capnometer to provide an index of changes in whole body metabolism and was maintained at 3.5–4.5% under basal conditions. *T*_BAT_ was measured from the left interscapular BAT pad, and postganglionic BAT SNA was recorded from the central cut end of a nerve bundle isolated from the right interscapular BAT pad.

BAT SNA was amplified (x 5,000–10,000) and filtered (1–300 Hz) by a CyberAmp 380 amplifier (Molecular Devices). All the physiological variables were digitized and recorded to a personal computer using Spike 2 software (version 7.10, CED, Cambridge, UK). For nanoinjection of drugs, a sharp glass pipette (tip inner diameter: 15–30 μm) filled with NMDA (*N*-methyl-D-aspartate, Sigma-Aldrich, M3262, 0.2 mM), C21 (200 µg/ml) or pyrogen-free 0.9% saline was perpendicularly inserted into the DMH and pressure-ejected (50 nl). All drugs were dissolved in pyrogen-free 0.9% saline. To identify the injection site, a small amount (∼ 5 nl) of 0.2% FluoSpheres (0.1 µm diameter, 0.2% solids in saline; F8801 or F8803, Thermo Fisher) was injected at the same site with the same pipette at the end of recordings.

For data analysis (Figure 6E), BAT SNA amplitudes were quantified using Spike2 in sequential 4-s bins as the square root of the total power (root mean square) in the 0–20 Hz band of the autospectra of each 4-s segment of the BAT SNA traces. The “power/4 s” traces (Figures 6C and 6D) were used for quantification and statistical analyses of changes in BAT SNA. Baseline values of BAT SNA, *T*_BAT_, expired CO_2_, HR and mean arterial pressure were the averages during the 1-min period immediately prior to NMDA injection. NMDA-evoked changes in *T*_BAT_, expired CO_2_, HR and mean arterial pressure were the differences between their baseline values and their peak values within 5 min (10 min for *T*_BAT_) after NMDA injection. NMDA-evoked changes in BAT SNA were the area under the curve (AUC) of the “power/4 s” trace above the baseline level (subtracted AUC) for 5 min after NMDA injection. NMDA-evoked increases in BAT SNA following C21 injection into the DMH were expressed as % of NMDA-evoked increases in BAT SNA following saline injection (Figure 6E).

### Anatomy and statistical analysis

We adopted the cytoarchitecture and nomenclature of most brain regions from those of Paxinos and Watson (2007). The raphe pallidus nucleus was nomenclaturally divided into the rostral (rRPa) and caudal (cRPa) parts at the rostral end of the inferior olivary complex (Nakamura et al., 2002, 2004).

Data are shown as the means ± SEM. Statistic comparison analyses were performed using a paired or unpaired *t*-test, an ordinary one-way ANOVA followed by Bonferroni’s multiple comparisons test, or a repeated measures two-way ANOVA followed by Bonferroni’s multiple comparisons test (Prism 9, GraphPad) as stated in the text and figure legends. All the statistic tests were two-sided. *P* < 0.05 was considered statistically significant.

## AUTHOR CONTRIBUTIONS

Y.N. and K.N. designed experiments. Y.N., T.Y. and K.N. performed experiments and analyzed and discussed data. Y.N., H.H. and K.N. generated and supplied *Ptger3-tTA* transgenic rats. N.K., H.H. and K.N. generated AAVs. Y.N. and K.N. wrote the manuscript, and all authors approved the manuscript.

## ACKNOWLEDGMENTS

We thank Akiya Watakabe for sharing the VGLUT2 antisense probe and Takeshi Kaneko for helpful discussion. This study was supported by the Funding Program for Next Generation World-Leading Researchers from the Japan Society for the Promotion of Science (LS070 to K.N.); Grants-in-Aid for Scientific Research (JP21K06767, JP17K08568, JP26860159 and JP23790271 to Y.N.; JP19K06954 and JP16K19006 to N.K.; JP21H02592 to H.H.; JP20H03418, JP16H06276 (AdAMS), JP16H05128,JP15H05932, JP26118508, JP26713009 and JP22689007 to K.N.) from the Ministry of Education, Culture, Sports, Science and Technology of Japan; the PRESTO program (JPMJPR13M9 to K.N.), the FOREST program (JPMJFR204D to H.H.) and Moonshot R&D (JPMJMS2024 to H.H. and JPMJMS2023 to K.N.) of the Japan Science and Technology Agency; the Japan Agency for Medical Research and Development (JP21wm0525002 to N.K., JP21dm0207112 to H.H. and JP21gm5010002 to K.N.); and by grants from the Hori Sciences and Arts Foundation (to Y.N.), Kato Memorial Bioscience Foundation, Foundation of Kinoshita Memorial Enterprise (to N.K.), Takeda Science Foundation, Nakajima Foundation, Uehara Memorial Foundation, Ono Medical Research Foundation, Brain Science Foundation, and Kowa Life Science Foundation (to K.N.). The maintenance of the *Ptger3-tTA* transgenic rat strain was supported by the National BioResource Project–Rat, Kyoto University (Kyoto, Japan).

## Notes

### Competing Interest Statement

The authors have declared no competing interest.

