## Supplemental Figures for "Prostaglandin EP3 receptor-expressing preoptic neurons bidirectionally control body temperature via tonic GABAergic signaling"

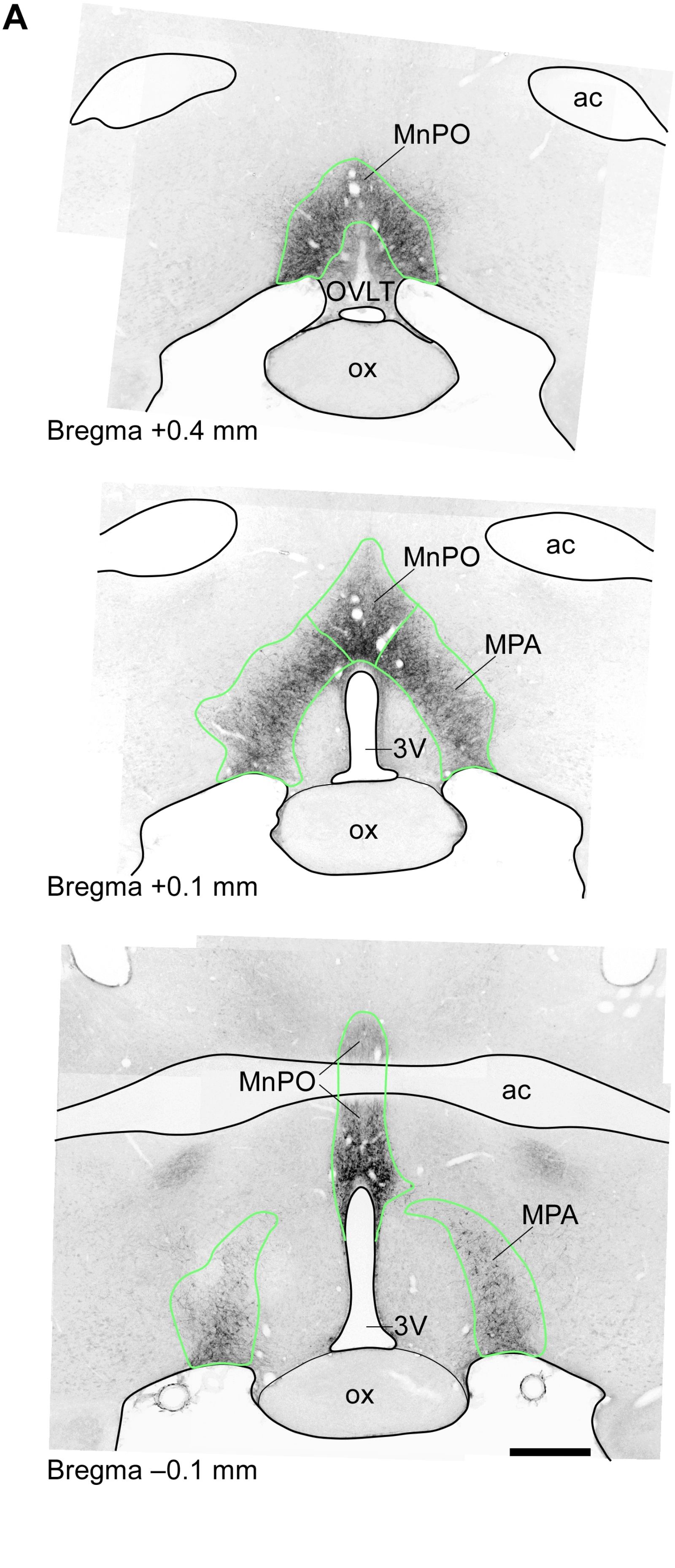
A
ac
MnPO
OVLT
ox
Bregma +0.4 mm
ac
MnPO
MPA
3V
ox
Bregma +0.1 mm
MnPO
ac
MPA
3V
ox
Bregma –0.1 mm

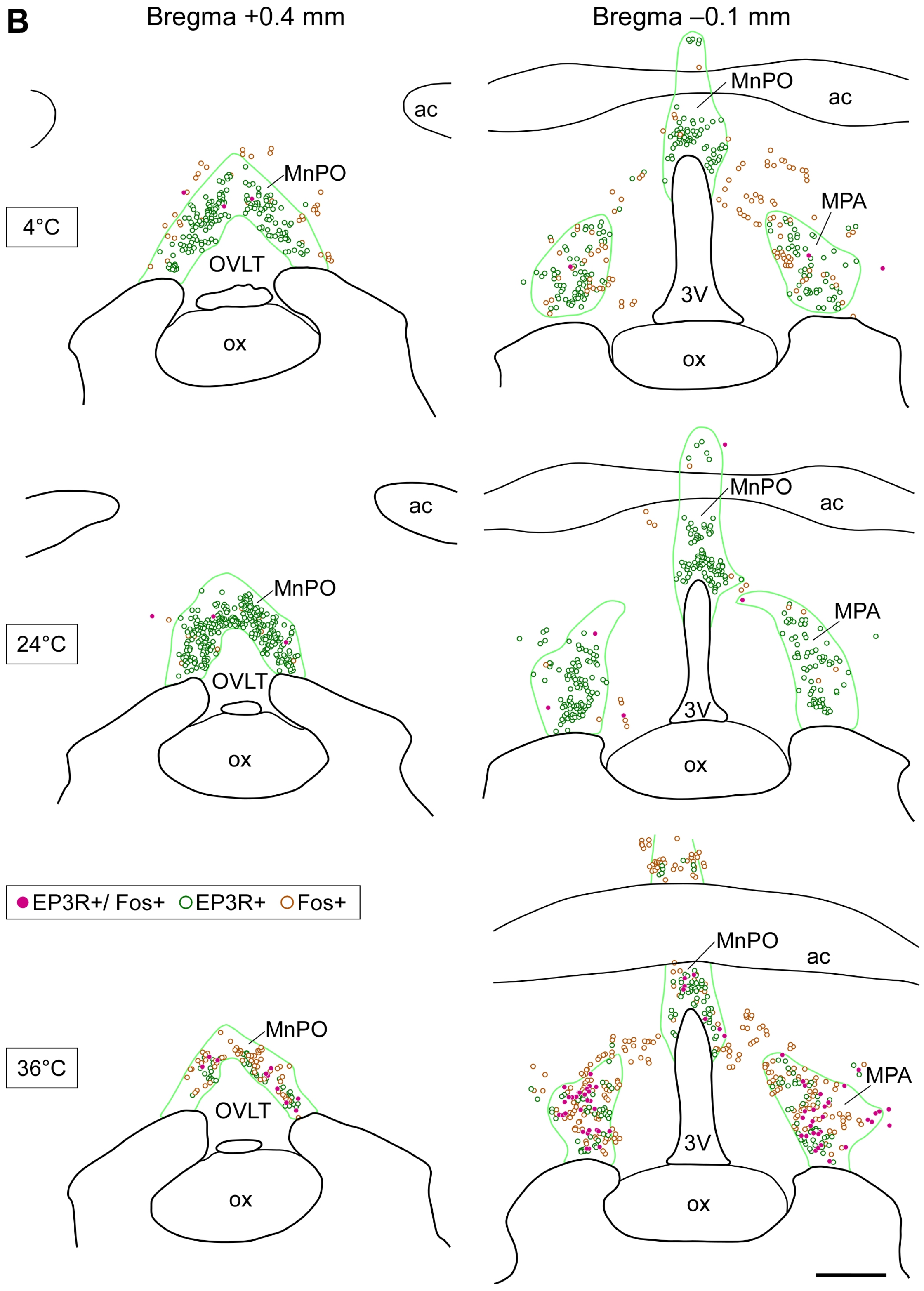
B
Bregma +0.4 mm
Bregma –0.1 mm
4°C
24°C
36°C
MnPO
OVLT
ox
ac
MPA
3V
EP3R+/ Fos+
EP3R+
Fos+

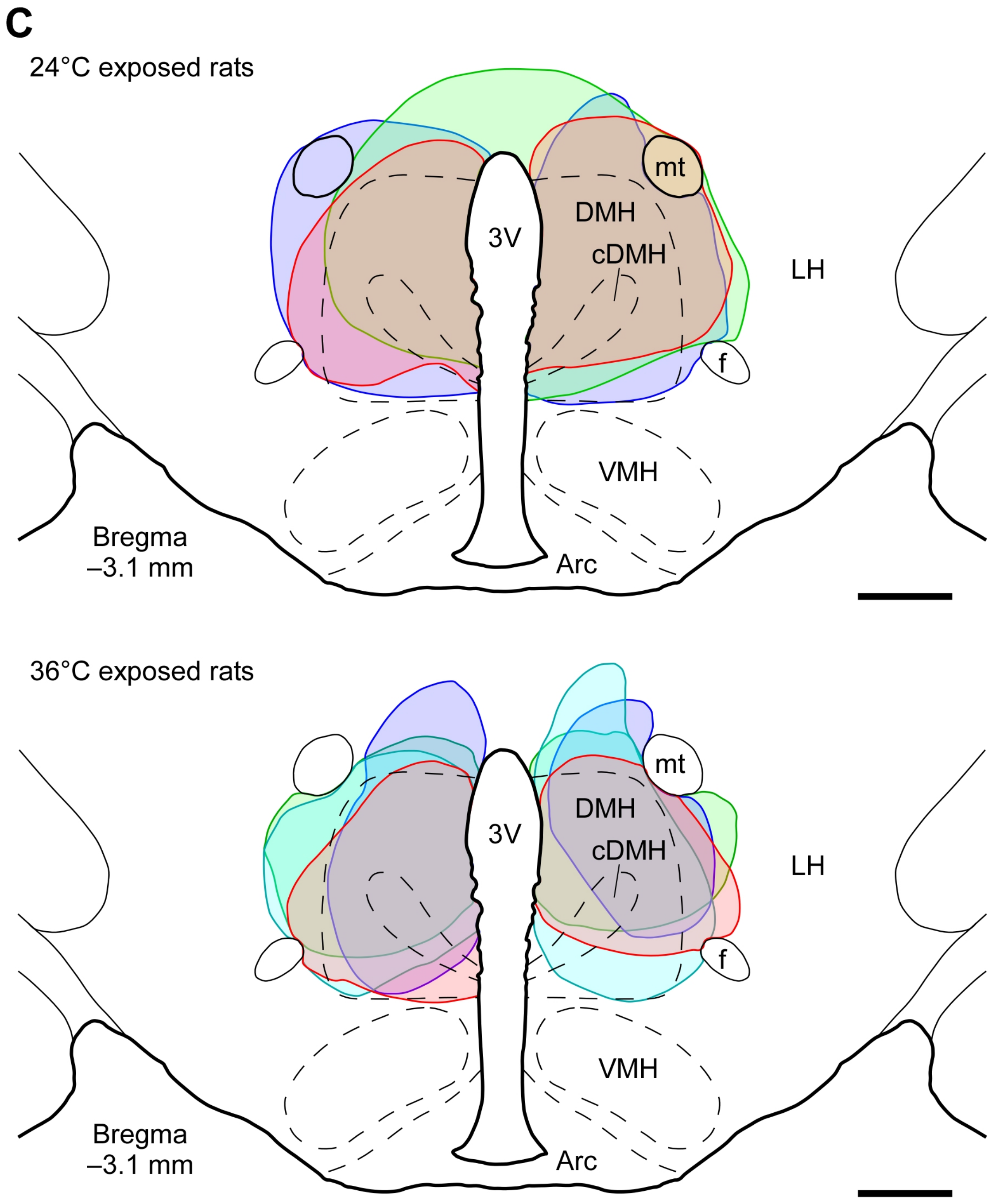
C
24°C exposed rats
mt
DMH
3V
cDMH
LH
f
VMH
Bregma
–3.1 mm
Arc
36°C exposed rats
mt
DMH
3V
cDMH
LH
f
VMH
Bregma
–3.1 mm
Arc

D

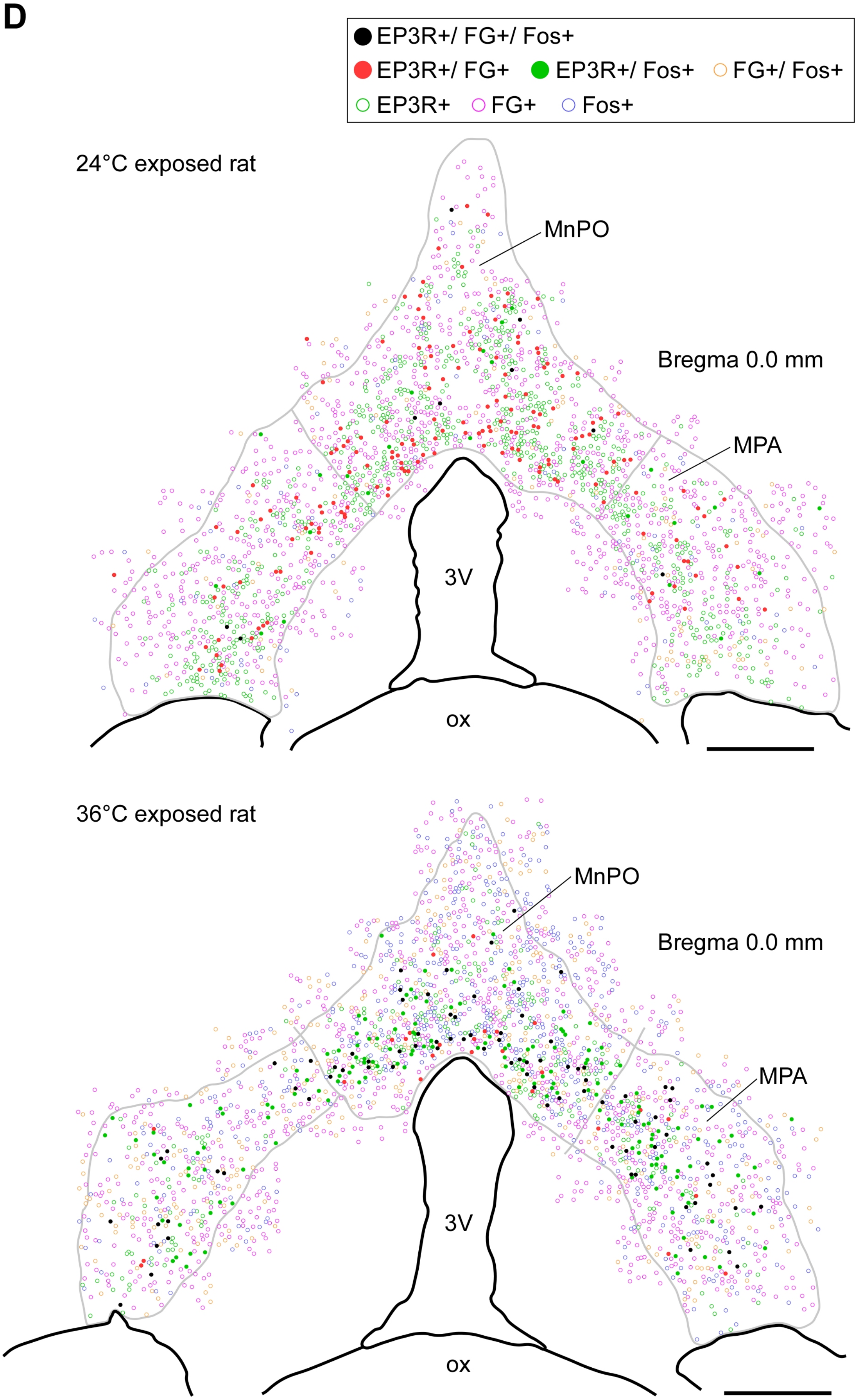
EP3R+/ FG+/ Fos+
EP3R+/ FG+
EP3R+/ Fos+
FG+/ Fos+
EP3R+
FG+
Fos+
24°C exposed rat
MnPO
Bregma 0.0 mm
MPA
3V
ox
36°C exposed rat
MnPO
Bregma 0.0 mm
MPA
3V
ox

**Figure S1**. **POA$^{EP3R}$ neurons are activated by ambient heat exposure and project to DMH, related to Figure 1**

(A) Composite immunofluorescence images that show the distribution of POA$^{EP3R}$ neurons. Green lines delineate the EP3R-immunoreactive areas in the MnPO and MPA. Scale bar, 0.5 mm. ac, anterior commissure; OVLT, organum vasculosum lamina terminalis.

(B) Representative distributions of POA cells with EP3R and/or Fos immunoreactivity following exposure to 4°C (cold), 24°C (control) or 36°C (heat). Green lines delineate EP3R-immunoreactive areas in the MnPO and MPA, as defined in (A). Scale bar, 0.5 mm. The distributions at the bregma +0.1 mm level in the same rats are shown in Figure 1C.

(C) Sites of bilateral FG injections in the DMH of the rats shown in Figure 1E. Scale bars, 0.5 mm. Arc, arcuate nucleus; cDMH, compact part of the dorsomedial hypothalamic nucleus; LH, lateral hypothalamic area; VMH, ventromedial hypothalamic nucleus.

(D) Representative distributions of POA cells labeled with FG, EP3R immunoreactivity and/or Fos immunoreactivity at the bregma 0.0 mm level following 2 hr exposure of rats to 24°C or 36°C ambient temperature. Gray lines delineate EP3R-immunoreactive areas in the MnPO and MPA. Scale bars, 0.3 mm.

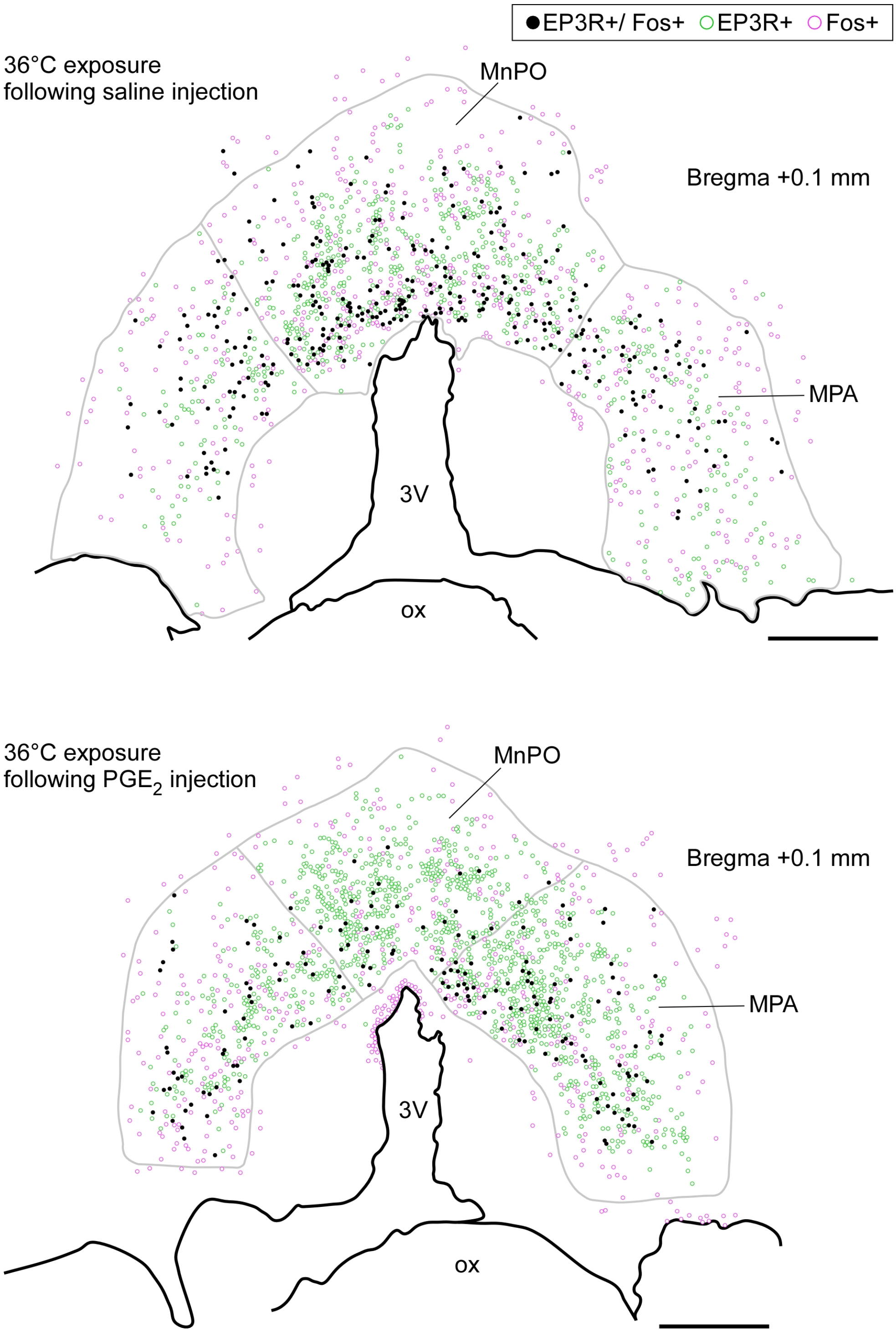


**Figure S2.  $PGE_2$ inhibits heat exposure-induced activation of $POA^{EP3R}$ neurons, related to Figure 2**

Representative distributions of POA cells with EP3R and/or Fos immunoreactivity at the bregma +0.1 mm level following 36°C ambient temperature after intracerebroventricular injection of saline or $PGE_2$.  Gray lines delineate EP3R-immunoreactive areas in the MnPO and MPA.  Scale bars, 0.3 mm.

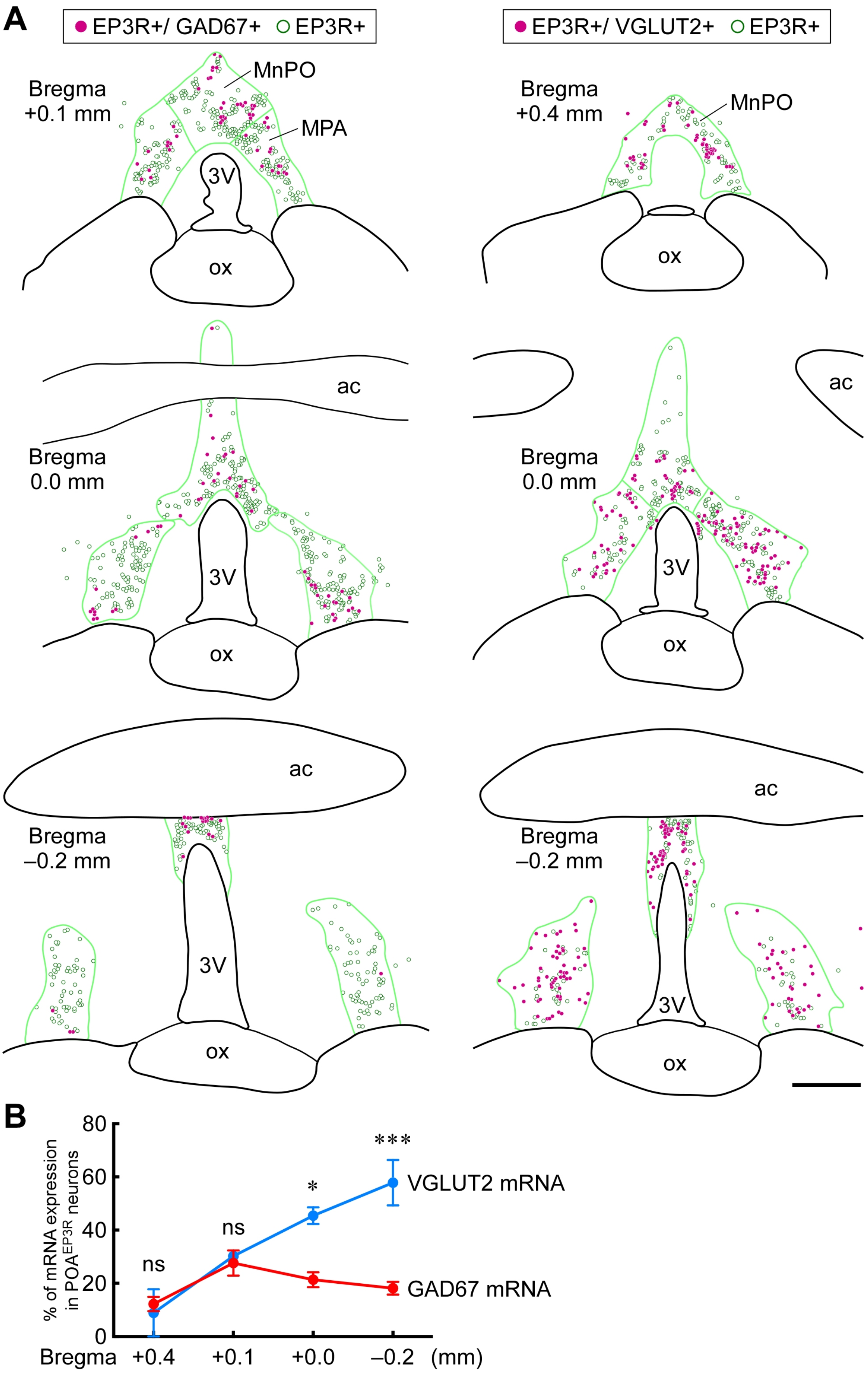
A
EP3R+/ GAD67+
EP3R+
EP3R+/ VGLUT2+
EP3R+
Bregma +0.1 mm
MnPO
MPA
3V
ox
Bregma +0.4 mm
MnPO
ox
ac
Bregma 0.0 mm
3V
ox
ac
Bregma 0.0 mm
3V
ox
ac
Bregma –0.2 mm
3V
ox
ac
Bregma –0.2 mm
3V
ox
B
% of mRNA expression in POA$^{EP3R}$ neurons
80
60
40
20
0
ns
ns
*
***
VGLUT2 mRNA
GAD67 mRNA
Bregma +0.4 +0.1 +0.0 –0.2 (mm)

C

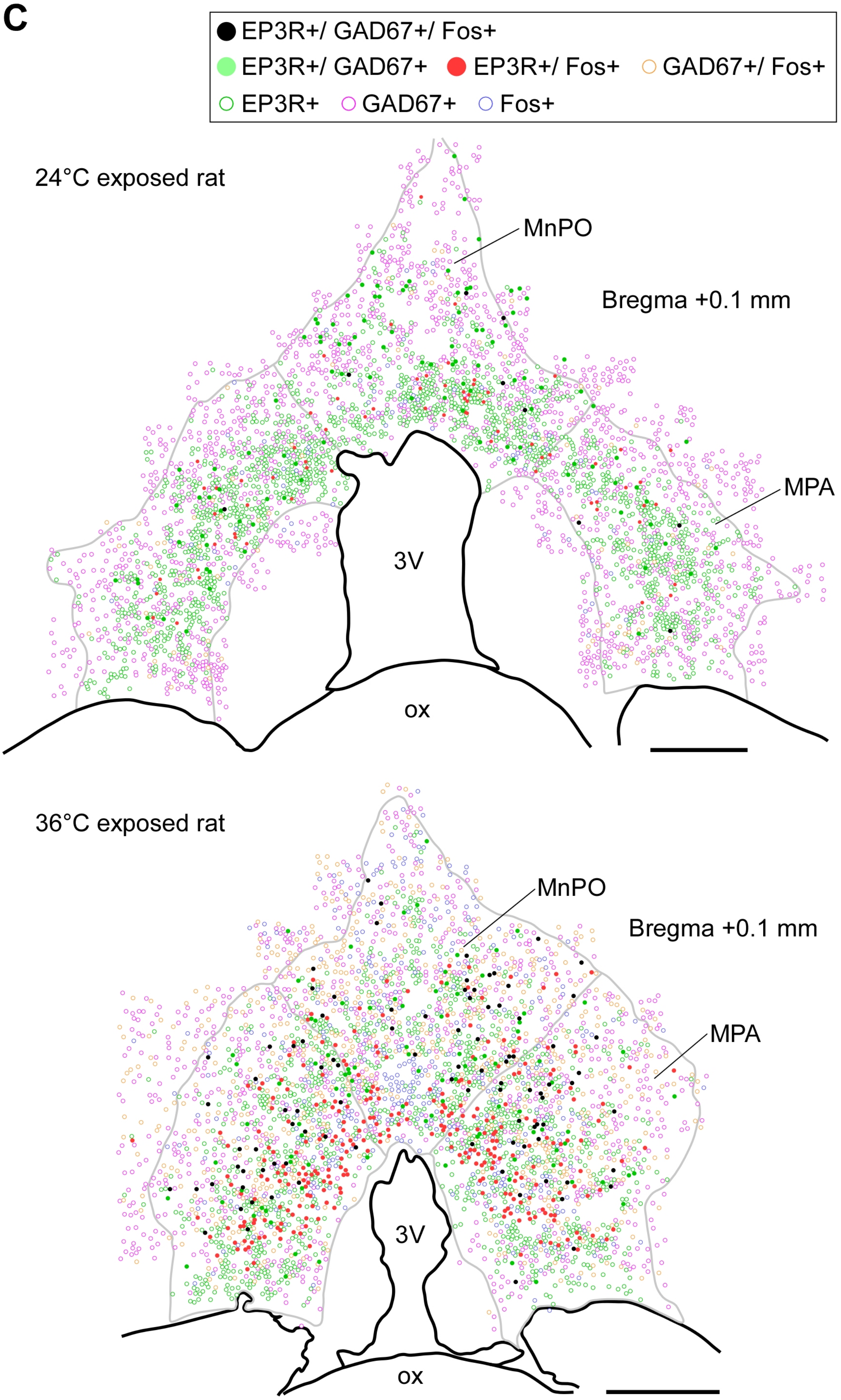
EP3R+/ GAD67+/ Fos+
EP3R+/ GAD67+
EP3R+/ Fos+
GAD67+/ Fos+
EP3R+
GAD67+
Fos+
24°C exposed rat
MnPO
Bregma +0.1 mm
MPA
3V
ox
36°C exposed rat
MnPO
Bregma +0.1 mm
MPA
3V
ox

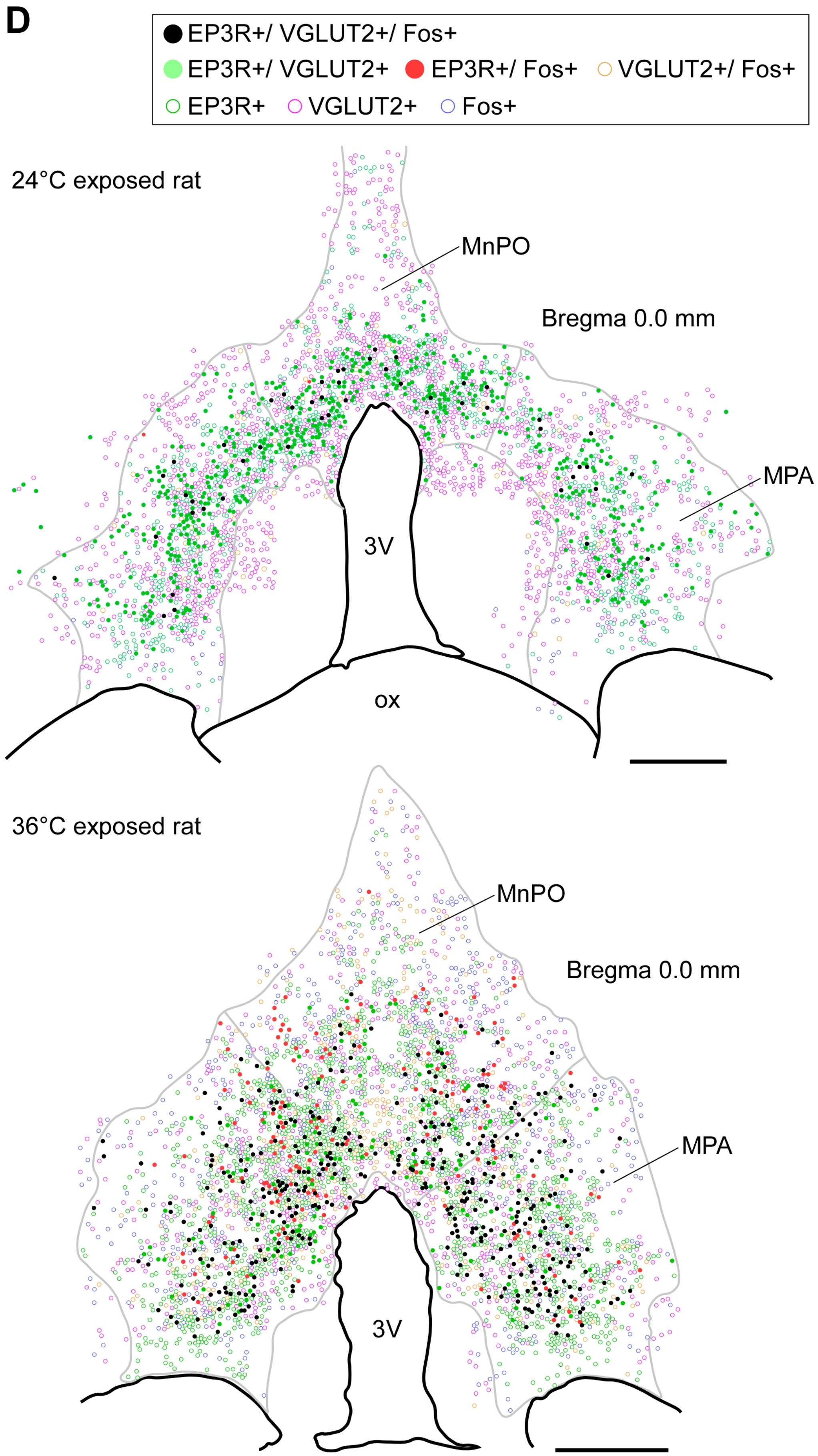
D
EP3R+/ VGLUT2+/ Fos+
EP3R+/ VGLUT2+
EP3R+/ Fos+
VGLUT2+/ Fos+
EP3R+
VGLUT2+
Fos+
24°C exposed rat
MnPO
Bregma 0.0 mm
MPA
3V
ox
36°C exposed rat
MnPO
Bregma 0.0 mm
MPA
3V

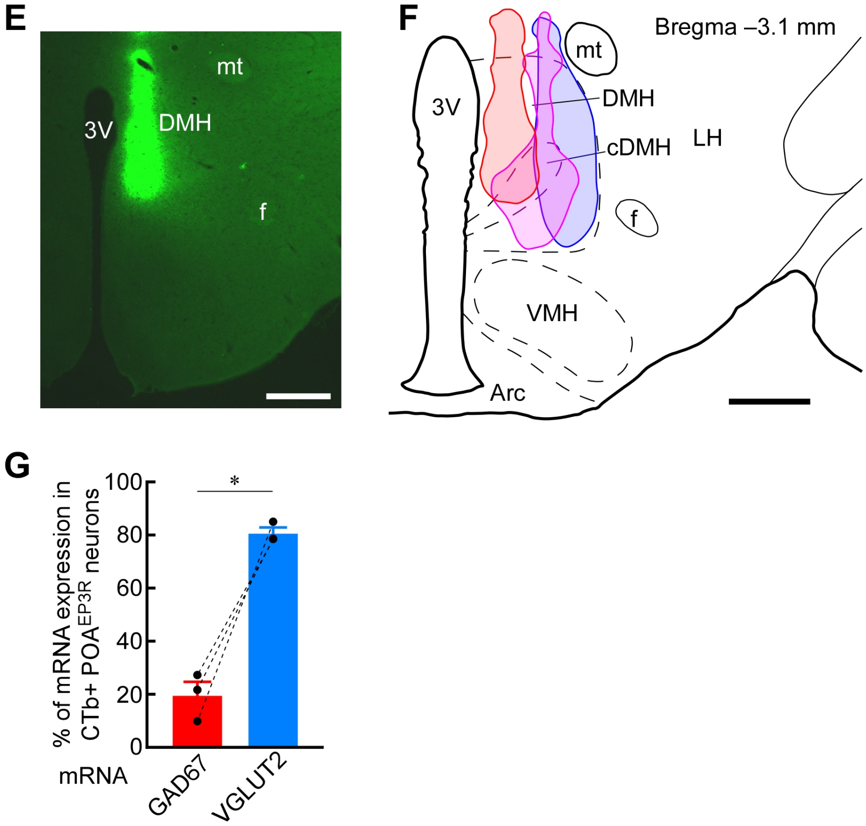


**Figure S3**. **POA$^{EP3R}$ neuronal cell bodies predominantly express glutamatergic marker rather than GABAergic marker, related to Figure 3**

(A) Representative distributions of POA$^{EP3R}$ neuronal cell bodies with and without GAD67 (left) or VGLUT2 (right) mRNA hybridization signals. Green lines delineate EP3R-immunoreactive areas in the MnPO and MPA. Scale bar, 0.5 mm.

(B) Percentages of GAD67 and VGLUT2 mRNA expression in POA$^{EP3R}$ neuronal cell bodies at different rostrocaudal levels in the POA. Data were analyzed by repeated measures two-way ANOVA (rostrocaudal level: $F_{3,18} = 21.65$, $P < 0.001$; mRNA: $F_{1,6} = 7.77$, $P < 0.05$; interaction: $F_{3,18} = 14.71$, $P < 0.001$) followed by Bonferroni's *post hoc* test (ns, not significant; *$P < 0.05$; ***$P < 0.001$, GAD67 vs VGLUT2). All values are means ± SEM.

(C and D) Representative distributions of POA cells with EP3R immunoreactivity, Fos immunoreactivity and/or GAD67 (C) or VGLUT2 (D) mRNA hybridization signals following 2 hr exposure of rats to 24°C or 36°C ambient temperature. Scale bars, 0.3 mm.

(E and F) A representative view of an injection into the DMH with Alexa488-conjugated CTb (E) and sites of unilateral CTb injections in the DMH (F) for all the rats shown in Figure 3H and (G). Scale bars, 0.5 mm.

(G) Percentages of GAD67 and VGLUT2 mRNA expression in CTb-labeled POA$^{EP3R}$ neuronal cell bodies ($n = 3$). *$P < 0.05$ (paired $t$-test; $t_2 = 8.49$). All values are means ± SEM.

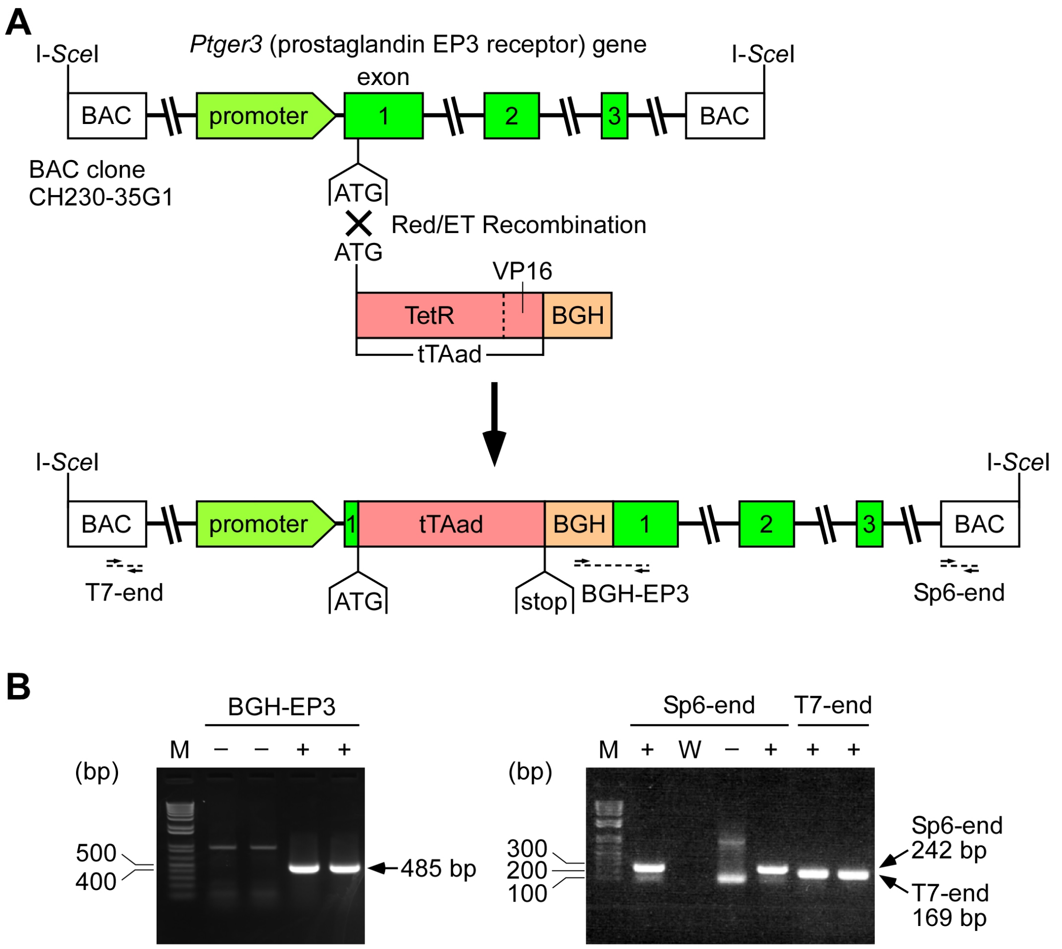


Primers

| | | |
|---|---|---|
| BGH-EP3 | Forward | 5'-**aaatgaggaaattgcatcgc**-3' |
| | Reverse | 5'-**cagtccgaacactgtcatgg**-3' |
| T7-end | Forward | 5'-**cgcgtaatacgactcactatag**-3' |
| | Reverse | 5'-**agtggtaacattcgtgtt**-3' |
| Sp6-end | Forward | 5'-**catacgatttaggtgacactatag**-3' |
| | Reverse | 5'-**tcaattcccagaacccacat**-3' |

**Figure S4. Production of *Ptger3*-tTA rats, related to Figures 4–6**

(A) DNA recombination to construct a *Ptger3*-tTA-BGH BAC. A tTAad-BGH cassette was inserted at immediately 3' to the start ATG codon of the *Ptger3* gene. Horizontal dashed lines indicate the PCR fragments that were detected for genotyping in (B).

(B) PCR genotyping. +, genomic DNA samples from *Ptger3*-tTA heterogygotes; –, genomic DNA samples from wild-type rats; M, DNA size marker; W, negative control with water instead of template genomic DNA.

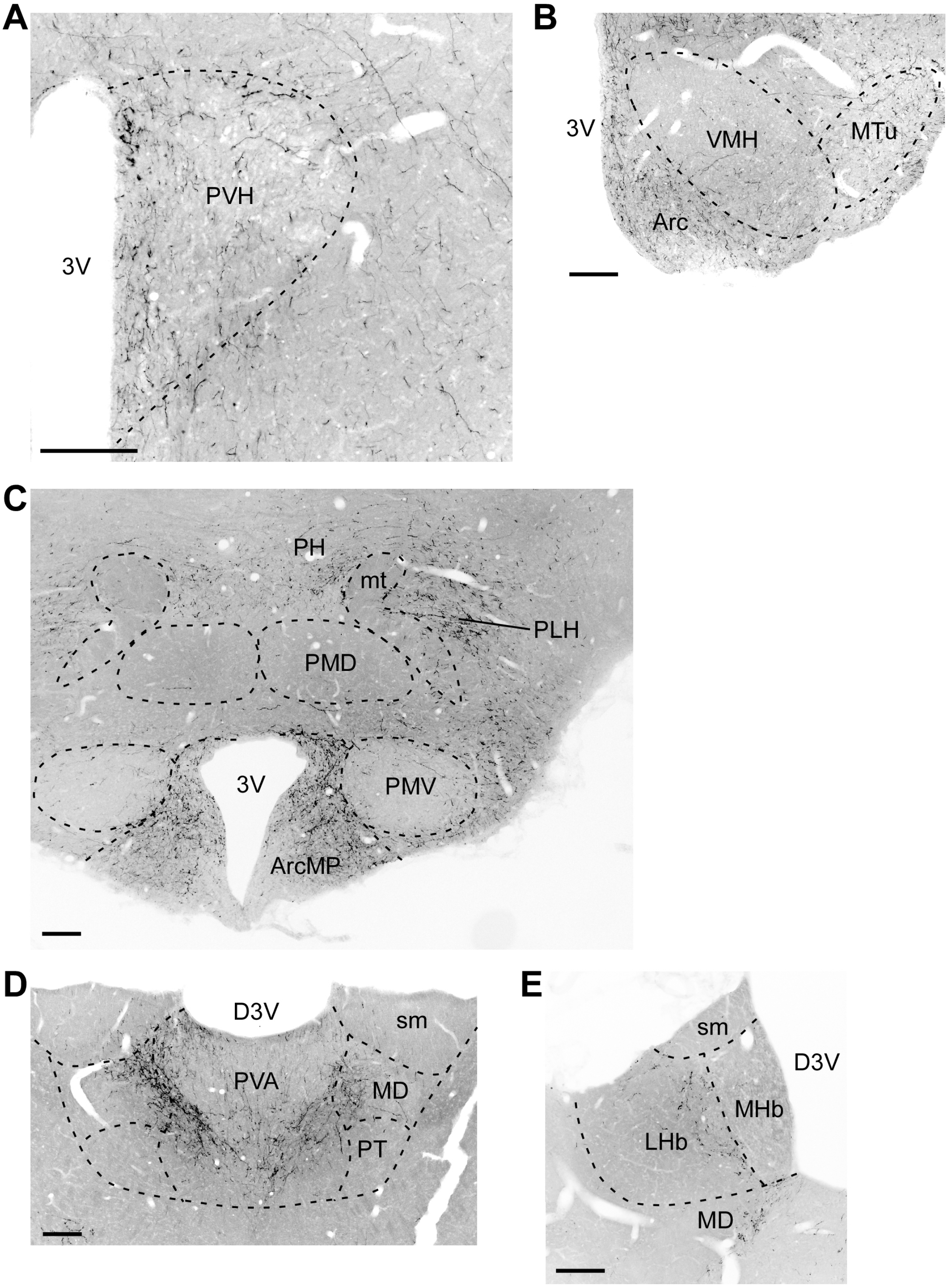
A
PVH
3V
B
3V
VMH
MTu
Arc
C
PH
mt
PLH
PMD
3V
PMV
ArcMP
D
D3V
sm
PVA
MD
PT
E
sm
D3V
MHb
LHb
MD

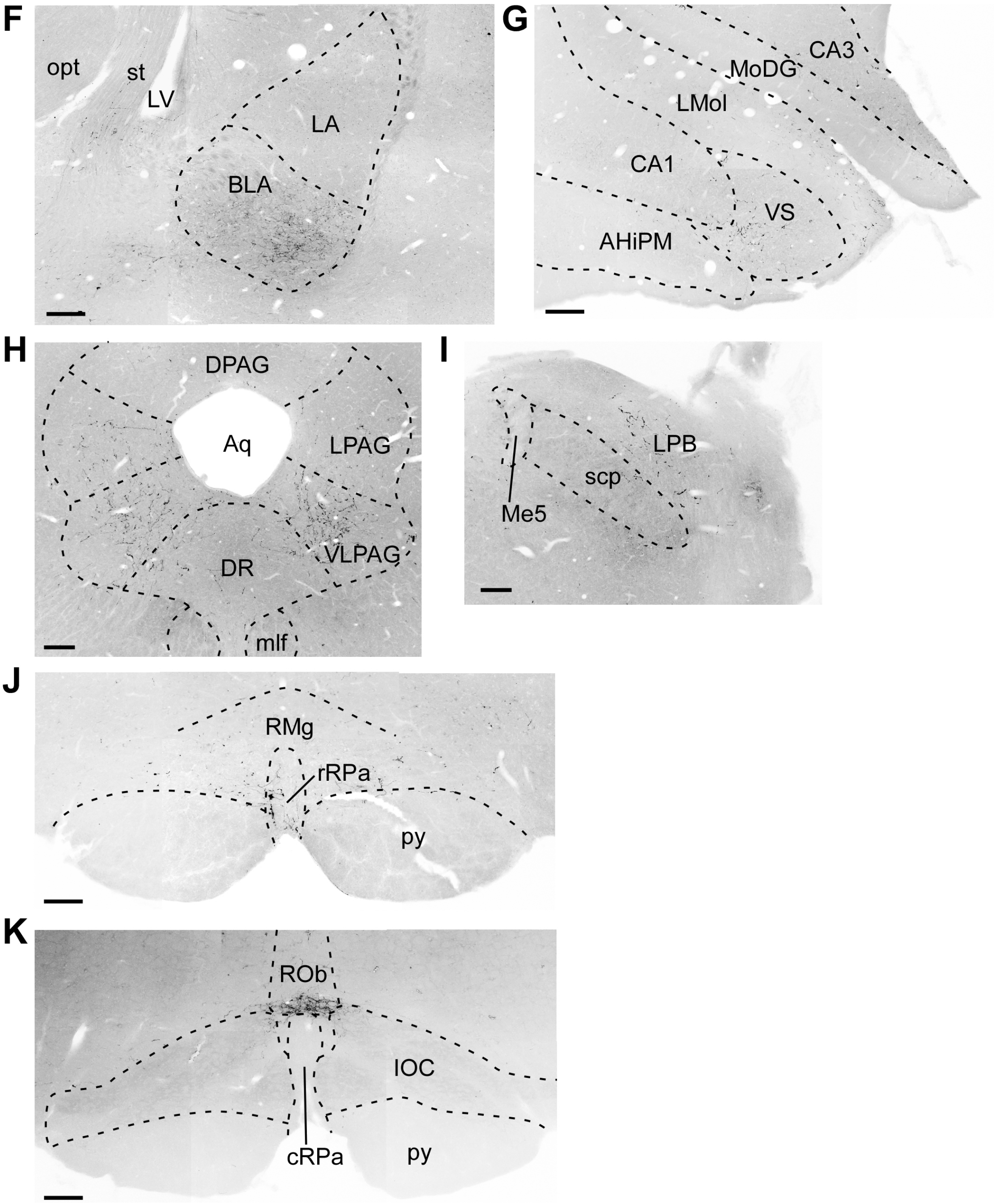


**Figure S5. Brain-wide distribution of axons from POA$^{EP3R}$ neurons, related to Figures 4C and 4D**

POA$^{EP3R}$ neurons in *Ptger3*-tTA rats were transduced with palGFP by injection of AAV-TRE-palGFP into the POA as shown in Figure 4A. Their axons throughout the brain were labeled with palGFP and further immunostained with a GFP antibody. Composite fluorescence images show the distributions of palGFP-labeled axons in the paraventricular hypothalamic nucleus (PVH) (A), ventral hypothalamus (B), mammillary body (C), anterior thalamus (D), habenula (E), amygdala (F), ventral hippocampus (G), periaqueductal gray (H), lateral parabrachial nucleus (LPB) (I), rostral ventromedial medulla (J) and caudal ventromedial medulla (K). Scale bars, 0.2 mm. AHiPM, posteromedial part of the amygdalohippocampal area; Aq, aqueduct; ArcMP, medial posterior part of the arcuate nucleus; BLA, basolateral amygdaloid nucleus; CA1, field CA1 of the hippocampus; CA3, field CA3 of the hippocampus; D3V, dorsal third ventricle; DPAG, dorsal periaqueductal gray; DR, dorsal raphe nucleus; IOC, inferior olivary complex; LA, lateral amygdaloid nucleus; LHb, lateral habenular nucleus; LMol, lacunosum moleculare layer of the hippocampus; LPAG, lateral periaqueductal gray; LPB, lateral parabrachial nucleus; MD, mediodorsal thalamic nucleus; Me5, mesencephalic trigeminal nucleus; MHb, medial habenular nucleus; mlf, medial longitudinal fasciculus; MoDG, molecular layer of the dentate gyrus; MTu, medial tuberal nucleus; opt, optic tract; PH, posterior hypothalamic nucleus; PLH, peduncular part of the lateral hypothalamus; PMD, dorsal part of the premammillary nucleus; PMV, ventral part of the premammillary nucleus; PT, paratenial thalamic nucleus; PVA, anterior part of the paraventricular thalamic nucleus; py, pyramidal tract; RMg, raphe magnus nucleus; ROb, raphe obscurus nucleus; scp, superior cerebellar peduncle; sm, stria medullaris of the thalamus; st, stria terminalis; VLPAG, ventrolateral periaqueductal gray; VS, ventral subiculum.

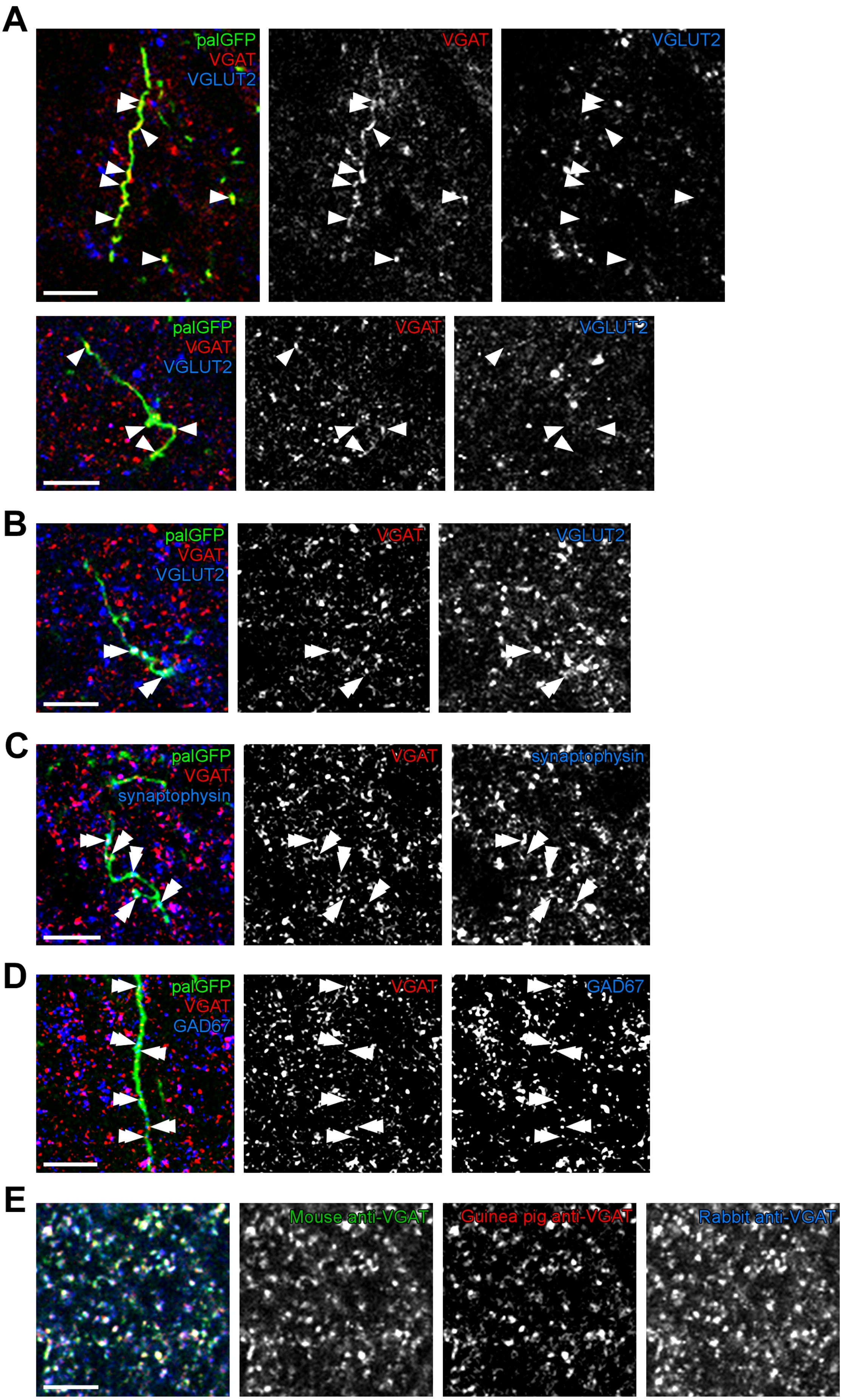
A
palGFP
VGAT
VGLUT2
VGAT
VGLUT2
palGFP
VGAT
VGLUT2
VGAT
VGLUT2
B
palGFP
VGAT
VGLUT2
VGAT
VGLUT2
C
palGFP
VGAT
synaptophysin
VGAT
synaptophysin
D
palGFP
VGAT
GAD67
VGAT
GAD67
E
Mouse anti-VGAT
Guinea pig anti-VGAT
Rabbit anti-VGAT

**Figure S6. POA$^{EP3R}$→DMH axon terminals predominantly contain GABAergic marker rather than glutamatergic marker, related to Figures 4E and 4F**

(A) Confocal images of palGFP-labeled (POA$^{EP3R}$ neuron-derived) axons with VGAT-immunoreactive puncta (arrowheads) in the DMH, additional to the example in Figure 4E. Scale bars, 10 μm.

(B) Confocal images of palGFP-labeled (POA$^{EP3R}$ neuron-derived) axons with VGAT- and VGLUT2-double immunoreactive puncta (double arrowheads) in the DMH. Scale bar, 10 μm.

(C) Confocal images of palGFP-labeled (POA$^{EP3R}$ neuron-derived) axons with VGAT- and synaptophysin-double immunoreactive puncta (double arrowheads) in the DMH. Scale bar, 10 μm.

(D) Confocal images of palGFP-labeled (POA$^{EP3R}$ neuron-derived) axons with VGAT-immunoreactive puncta accompanied by GAD67 immunoreactivity (double arrowheads) in the DMH. Scale bar, 10 μm.

(E) Confocal images showing triple immunofluorescence staining in the DMH with mouse, guinea pig and rabbit antibodies to VGAT. The merged image (left end) shows overlapped distributions of immunoreactivities of the three antibodies. Scale bar, 10 μm.

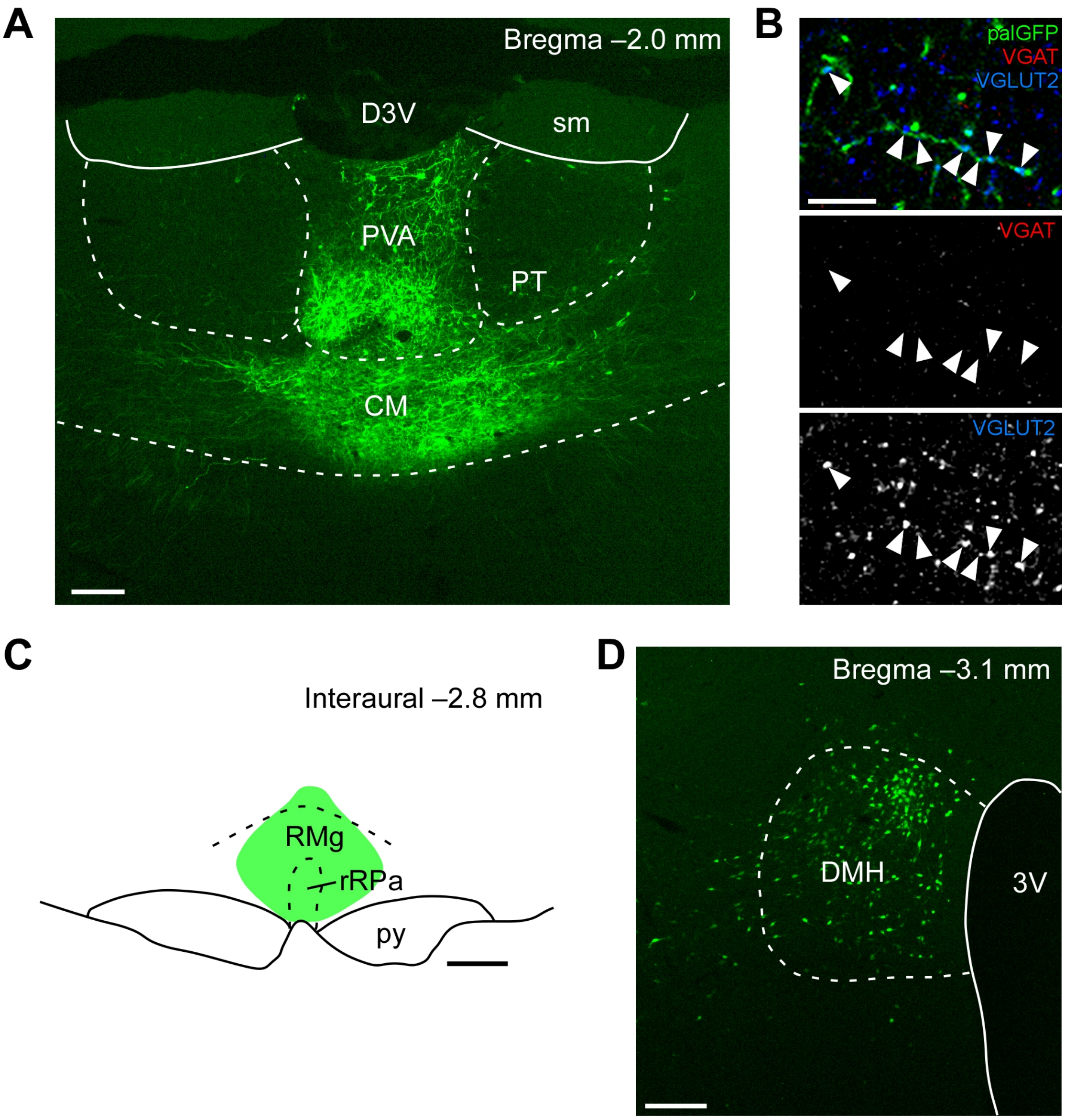


**Figure S7. Thalamocortical axons contain abundant VGLUT2-immunoreactive puncta and DMH contains densely localized neurons projecting to the rMR, related to Figures 4E, 4F and 4H**

(A and B) EP3R-expressing thalamic neurons were transduced with palGFP by injection of AAV-TRE-palGFP into the anterior midline thalamus of *Ptger3*-tTA rats (A). Confocal images (B) show that their palGFP-labeled axons in the area 2 of the cingulate cortex contained abundant VGLUT2-immunopositive puncta (arrowheads). Scale bars, 0.2 mm (A) and 10 µm (B). CM, central medial thalamic nucleus.

(C and D) Retrograde labeling of DMH→rMR projection neurons. An injection of FG into the rMR (green area in C) resulted in retrograde FG-labeling of many DMH neurons, which were densely distributed particularly in the dorsal part of the DMH. Scale bars, 0.5 mm (C) and 0.2 mm (D).
